# Src-Family B Kinase Inhibition Promotes Lung Endothelial Dysfunction and Guides Drug Discovery

**DOI:** 10.1101/2021.09.27.462034

**Authors:** Adam M. Andruska, Xuefei Tian, Md Khadem Ali, Edda Spiekerkoetter

## Abstract

Protein tyrosine kinase (PTK) inhibition is efficacious in treating conditions ranging from cancer to fibrosis but can be limited by endothelial cell dysfunction. In trials of protein tyrosine kinase inhibitors (TKIs) a broad range of vascular effects is observed, inducing clinically detrimental endothelial cell apoptosis, impaired barrier function, or improved pulmonary vascular resistance in pulmonary arterial hypertension (PAH). We hypothesize this range of effects is due to subsets of PTKs either impairing or promoting endothelial homeostasis by modulating Bone Morphogenetic Protein receptor 2 (BMPR2) signaling, a pathway essential for vascular development and dysfunctional in PAH. In a high-throughput siRNA screen we find SRC-Family B PTKs activate whereas SRC-Family A PTKs suppress BMPR2 signaling, measured by the transcription factor inhibitor of differentiation 1 (Id1). Induced loss of function of the strongest ID1 activating PTK LCK (a Src-B kinase) in human pulmonary artery endothelial cells suppresses BMPR2 signaling and induces multiple measures of endothelial dysfunction. However, loss of function of the strongest ID1 inhibitor PTK FYN (a Src- A kinase) does the opposite. Whole-genome transcriptional analysis identifies two multi-gene signatures inversely regulated by LCK and FYN we term “endothelial” and “inflammatory”. To find TKIs mimicking selective Src-A and Src-B inhibition, we use Connectivity map to identify drugs connecting to the endothelial and inflammatory signatures. We find several TKIs: a pro-inflammatory (Regorafanib), a BMPR2 potentiating (Brivanib), and a BMPR2 suppressing (Quizartinib). Here we show a dichotomy in pathway regulation by Src- A and -B kinases that may have utility in transcriptionally based drug discovery.

## Introduction

Cells have evolved protein kinases to facilitate signal transduction. 518 protein kinases are present in humans of which 90 are Protein Tyrosine Kinases (PTKs). PTKs subdivide into receptor (e.g. VEGFR) and non-receptor, “cytoplasmic” (e.g. SRC) groups [1]. Receptor-PTKs enable intercellular ligand-based communication whereas non-receptor PTKs amplify external and internal signals and control functions like proliferation and motility. Many non-receptor PTKs are oncogenes (e.g. BCR-ABL) which has driven the development of Tyrosine Kinase Inhibitors (TKIs). TKIs are now approved for conditions ranging from cancer to pulmonary fibrosis [2]. Though successful in treating disease, TKIs have mixed effects in pulmonary vascular biology.

Pulmonary Arterial Hypertension (PAH) is a progressive and fatal obliterative vasculopathy of the pulmonary arterioles characterized by endothelial cell dysfunction (ECD) and apoptosis in concert with smooth muscle cell proliferation [3–6]. Though limited by adverse reactions, the TKI Imatinib improves clinical PAH likely by acting on the receptor PTKs PDGFR and c-KIT [7, 8]. Paradoxically, TKIs like Dasatinib [9–11], Bosutinib [12], Ponatinib [13], and Lorlatinib [14] are either known causes of PAH or are implicated in PAH development. Dasatinib may achieve this by acting on additional PTKs outside of PDGFR and c-KIT [10, 15, 16], secondary induction of reactive oxygen species (ROS) in pulmonary artery endothelial cells (PAECs) [11], or impairing endothelial cell (EC) barrier permeability in a Rho-kinase dependent manner [17, 18]. In support of Dasatinib acting on a broader set of PTKs, a recent pharmacovigilance study showed the SRC family PTKs LCK, LYN, and YES are disproportionately associated with the development of PAH [19].

Despite association between PTKs, TKIs, and PAH development, knowledge of how PTKs act on specific cell types and signaling pathways continues to evolve. Since it is observed that (1) different TKIs generate markedly different clinical phenotypes and (2) specific PTKs associate with human disease [19], we theorized that individual PTKs may disproportionately target signaling pathways required for pulmonary vascular integrity.

One such pathway is the Bone Morphogenetic Protein Receptor 2 (BMPR2) pathway. Heterozygous mutations in *BMPR2* are the most common cause of hereditary PAH [4, 20]. *In vitro* silencing of *BMPR2* recapitulates many features of ECD [21]. However, because *BMPR2* disease penetrance is 20%, other genetic and environmental factors are implicated in PAH development [4]. Work investigating the interaction of PTKs with BMPR2 signaling shows inhibition of the non-receptor PTK SRC can improve BMPR2 signaling by increasing BMPR2 receptor plasma membrane localization [22, 23]. Conversely, we recently identified that the Src-family tyrosine kinase LCK has the potential to support BMPR2 signaling [24]. Given the opposite effects on BMPR2 signaling by LCK and SRC, we wondered if subsets of PTKs may differentially affect BMPR2 signaling and EC health.

To test this, we examined a broad group of PTKs in a high-throughput siRNA screen and found that Src-family non-receptor PTKs divide into BMPR2-supportive and -repressive groups when using a canonical BMPR2 responsive transcription factor, Inhibitor of Differentiation 1 (ID1) as a readout for BMPR2 signaling. Surprisingly, Src family PTKs modulate BMPR2 in a way that respects an evolutionarily defined boundary separating Src Family A (SrcA) and Src Family B (SrcB) PTKs. Within these groups, we found that the SrcB LCK is the most BMPR2-activating kinase while the SrcA FYN is the most BMPR2-repressive. We also find this divided effect on cell signaling pathways extends to canonical NF-κB signaling. Using the transcriptomic signature of SrcA FYN and SrcB LCK knockout in PAECs, we use the publicly available Connectivity Map (CMap) database to find drugs that mirror the shared effects of *FYN* and *LCK* knockout. This led us to two TKIs, Brivanib and Quizartinib, that potentiate and suppress canonical BMPR2 signaling in PAECs. We demonstrate for the first time that the evolutionary divide between SrcA and SrcB can extend to the differential control of cell signaling pathways, with SRC PTKs split into two groups: a pro-BMPR2 “endothelial” SrcB, and an anti-BMPR2 “inflammatory” SrcA. This may further guide the selection of TKIs that achieve therapeutic goals and avoid ECD in diseases such as PAH.

## Results

### A High-Throughput screen finds that Src-Family A PTKs are BMPR2-Repressive, and Src-Family B PTKs are BMPR2-Activating

To assess canonical BMPR2 signaling, we used *Id1* as a transcription factor read-out for BMPR2 activation. As previously described, we employed a C2C12 mouse myoblastoma cell line stably transfected with a BMP response element Luciferase Id1 (BRE-Luc-Id1) reporter [25]. Cells were systematically transfected with a murine-wide siRNA library targeting 22,124 genes and then treated with BMP4 in the Stanford High-Throughput Bioscience Center. BMPR2 activity by luciferase expression was quantified by luminescence and cell viability was assessed with a tryptan-blue stain (Figure 1A). The mean change in *Id1* expression and cell viability in response to knockout of one of the 22,124 genes across three replicates was calculated and is presented in aggregate in Figure 1B. Genes to the left of 0 on the x-axis reduce and genes to the right of 0 on the x-axis increase *Id1* expression when targeted by siRNA. Using published phylogenetic classifications of receptor- and non-receptor-PTKs [1], representative PTKs from major families were chosen and highlighted within the high throughput screen (HTS). An even subdivision of PTKs activated (20 of 41) and repressed (21 of 41) *Id1* expression (Figure 1B, red dots). The PTKs with the greatest positive and negative changes, respectively, in Id1 transcription were the non-receptor PTKs Lck and Fyn, both members of the Src-Family of protein tyrosine kinases (Figure 1C). Focusing specifically on the Src-Family, we found the magnitude and direction of change in *Id1* expression induced by PTK knockout matched exactly the phylogenetic division between SrcA and SrcB (Figure 1D and 1E). In summary, our agnostic, high throughput screen shows that Id1 modulation by Src-family kinases respects the phylogenetic division between SrcA and SrcB.

**Figure 1:**
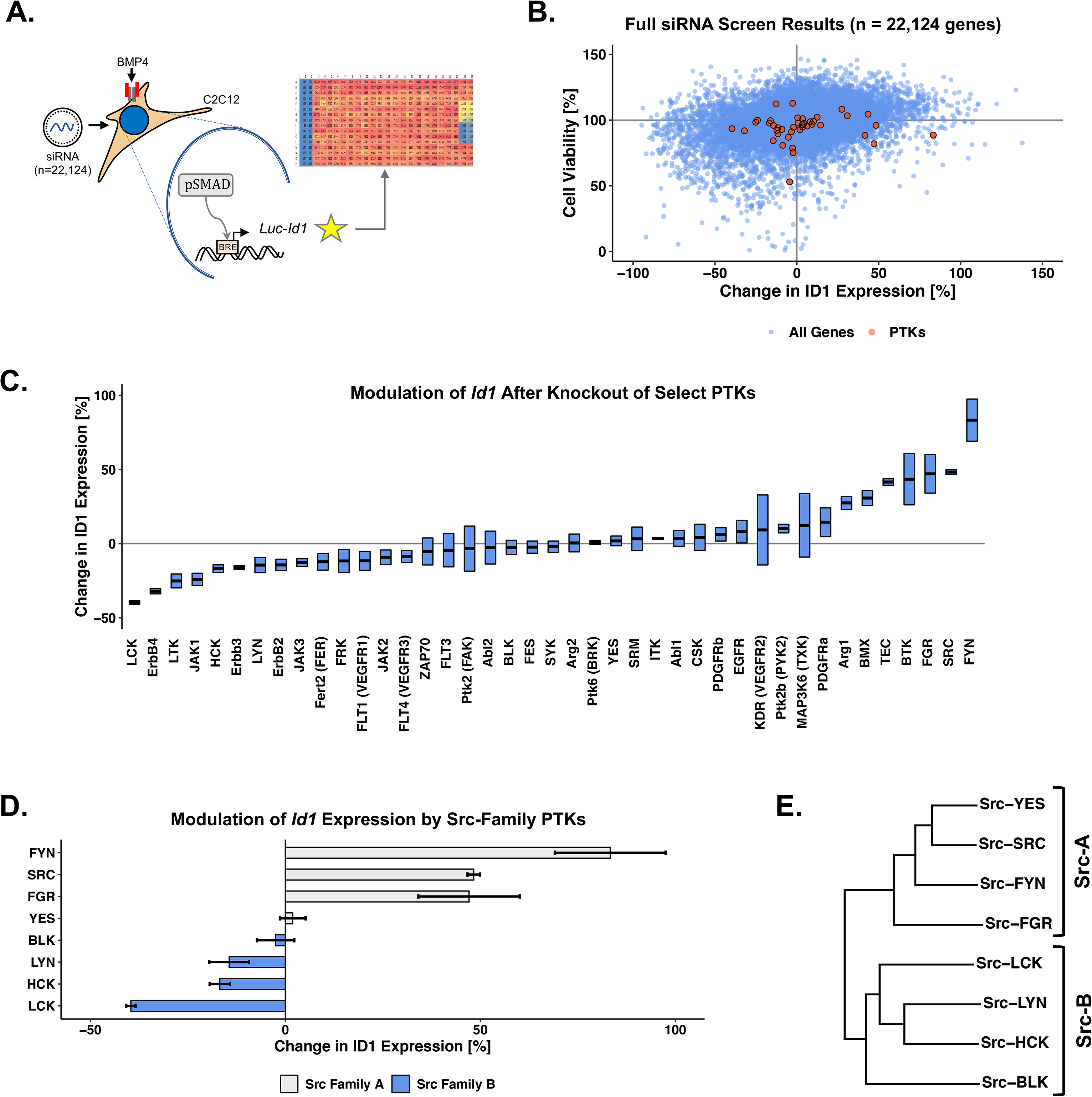
A High-Throughput Screen (HTS) using a C2C12 mouse myoblastoma Id1-Luciferase Reporter cell line finds Src-Family A PTKs are inherently Bmpr2-Repressive, and Src-Family B PTKs are inherently Bmpr2-Activating. (A) HTS treatment and assessment strategy. (B) Change in Id1-linked luciferin expression and cell viability after gene knockout across 22,124 gene. Red dots = major PTK families. (C) Change in Id1-linked luciferin expression in selected receptor and non-receptor PTKs. Bars = mean ± standard deviation (SD). Fyn and Lck generate the greatest change in Id1 expression. (D) Change in Id1-linked luciferin expression for Src-Family PTKs. Bars: mean ± SD . Overall magnitude of Id1 change correlates to the PTK’s sub-family (shown in (E), reconstructed from data in [1]): Src-A PTK knockout increases and Src-B PTK knockout decreases Id1 expression.

### LCK mRNA and protein are found in endothelial cells of human pulmonary arteries and correlate with ID1 expression both in health and in disease

We next aimed to find a human cell lineage suitable to study the relationship between SrcA, SrcB and BMPR2 pathway members, focusing on FYN as the representative SrcA kinase and LCK as the representative SrcB as they generated the greatest magnitude of change in our HTS. Dogmatically, SrcA are widely expressed while SrcB are restricted to specific lineages [1]. FYN has been studied in ECs and smooth muscle cells [26, 27] while LCK has traditionally been described in T-Cells and NK-Cells. Recently we and others have shown LCK is expressed in cultured human PAECs and HUVECs [24, 28], though where LCK is expressed in intact lung tissue and if its expression is altered in disease remained unknown. Single molecule RNA *in situ* hybridization (sm-FISH) found low *LCK* mRNA expression in the endothelium (white arrows) compared to higher expression in sparse ACTA2/CDH5 negative cells (yellow arrows) presumed to be lymphocytes (Figure 2A). LCK protein was identified in cultured human PAECs with perinuclear localization (Figure 2B). Human PAECs obtained from both PAH patients and donor controls (Patient Characteristics can be found in Supplemental Table 1) expressed LCK protein, but in a heterogeneous fashion (Figure 2C). Assessing for ID1 and SNAIL/SLUG expression as surrogates for active BMP and TGF-β signaling in these cell lines found a significant correlation between ID1 and LCK in both control and PAH patient samples (r = 0.68, p = 0.043, Figure 2D). The expression level of *LCK*, *FYN* and BMPR2 pathway member genes were quantitated lymphocytes from healthy donor peripheral blood mononuclear cells (PBMC) undergoing single cell RNA sequencing (publicly available, 10x Genomics and [29]). High *LCK* expression was verified in T-Cells and NK-Cells (Figure 2E, Sup. Figure 1). However, *BMPR2* and *ID1* were expressed in few PBMC cells (<20%), with low expression in all cell lineages save for dendritic cells. With this confirmation that LCK, ID1 and BMPR2 co-expression was poor in lymphocytes, we focused our attention on PAECs in the human lung where co-expression was confirmed.

**Figure 2:**
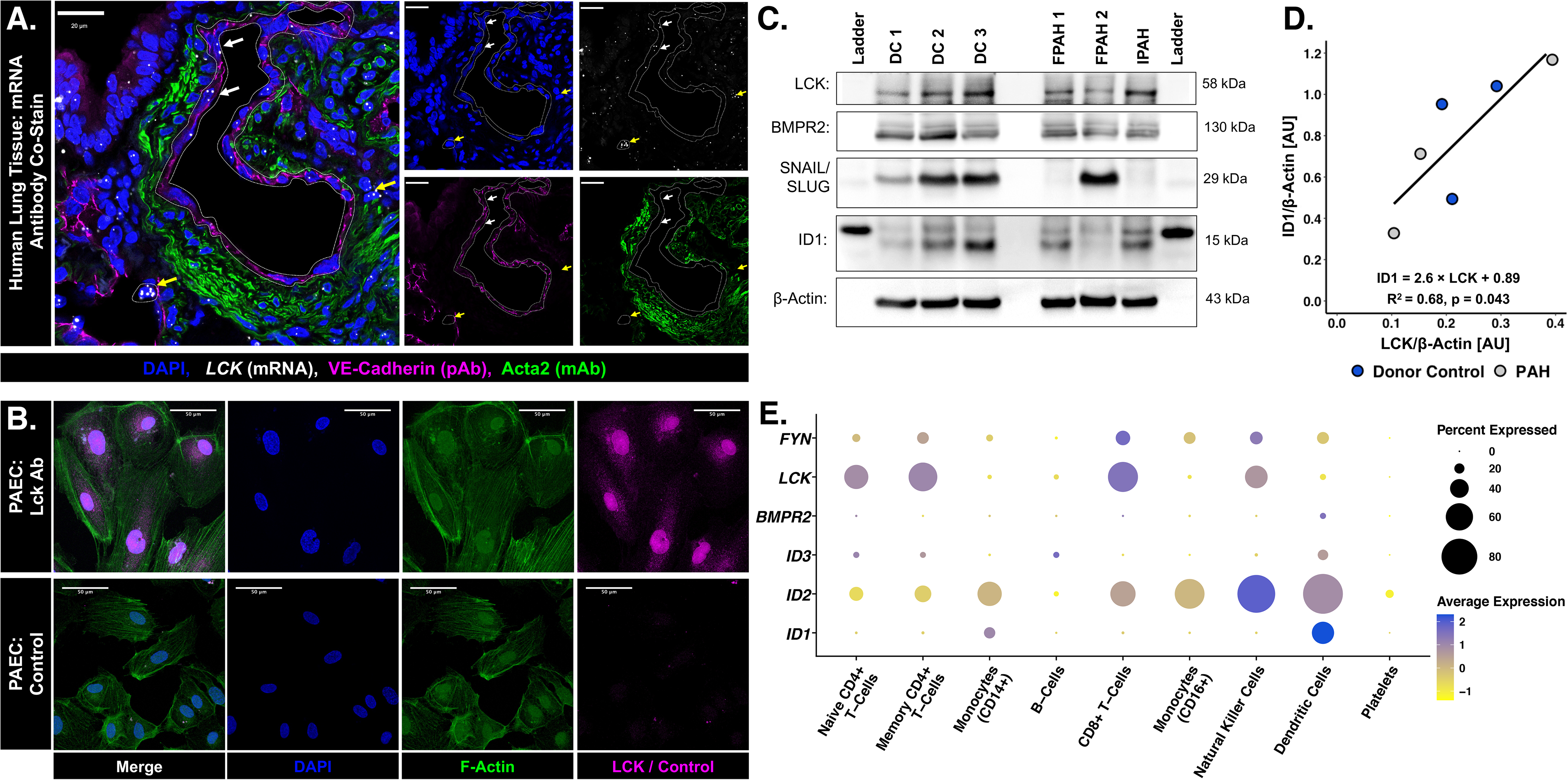
LCK is expressed in the endothelium of PAECs and correlates with ID1 expression. SC- RNAseq of human PBMCs suggests *BMPR2* and *ID1* is low in lymphocytes that have high expression of *LCK* or *FYN*. **(A)** smFISH in human lung for *LCK* mRNA (white), VE-Cadherin (magenta, EC Marker), ACTA2 (green, Smooth Muscle Marker), and DAPI (blue, nuclear marker). LCK/VE-Cadherin colocalizes in pulmonary arteries (white arrows). *LCK*-Hi, VE-Cadherin/ACTA2 negative cells (yellow arrow) likely represent lymphocytes. Bars = 20 μm **(B)** Peri-nuclear LCK expression in human PAECs. Bar = 50 μm. **(C-D)** Immunoblotting of protein lysate from human PAECs from explanted lungs of donor controls and PAH patients showing correlation between LCK and ID1 expression. **(E)** SC-RNAseq analysis of 2700 PBMCs from a healthy donor (full analysis Sup Figure 1) using the Seurat SC-RNA seq pipeline [60]. *LCK* and *FYN* are highly expressed in CD4+ and CD8+ T-Cells, and NK Cells. However, *BMPR2* and *ID1* are highly expressed in Dendritic Cells. *ID2* rather than *ID1* or *ID3* is highly expressed in lymphocytes, NK cells, and dendritic cells. Dot size indicates percent of cells expressing gene of interest (cutoff of >1 count per cell). Average Expression is log-scaled Z-score of counts for each gene in each cluster.

### Silencing *Lck* significantly impairs both basal and BMP9-mediated canonical BMPR2 signaling whereas silencing *Fyn* does the opposite

We next aimed to assess if the opposing effects of LCK and FYN on ID1 expression held true in PAECs. Because an interaction between the SH3 regions of LCK and SMAD proteins has been demonstrated [30], we extended our assessment to the canonical BMPR2/TGF-β second messenger SMAD proteins. SrcB LCK silencing (siLCK) in PAECs, confirmed by *LCK* mRNA (Figure 3A) and LCK protein (Figure 3B), reduced *ID1* mRNA expression and increased *SNAI1* expression, a read-out for activated TGF-β signaling. Furthermore, siLCK significantly impaired BMP9-induced ID1 expression (Figure 3C, and Sup. Figure 2A,B) and reduced SMAD1 phosphorylation, though to a lesser degree. PSMAD3 levels mildly, but not significantly, increased with either siLCK or TGF-β1 stimulation. When the SrcA *FYN* was silenced (siFyn) in PAECs *ID1* and *BMPR2* mRNA expression significantly increased at 48 hours (Figure 3D). Notably, the increase in ID1 expression following BMP9 stimulation was further augmented by *FYN* knockout. (Figure 3E), though we could not detect a change in pSMAD1 expression in BMP9 treated PAECS with or without *FYN* silencing, perhaps due to a saturation effect. BMP-9 treated PAECs had reduced pSMAD3 expression after *FYN* silencing relative to nontargeted controls (Figure 3E). Taken together, we confirmed that LCK and FYN have opposing effects in the pulmonary endothelium with respect to canonical BMPR2 signaling. *LCK* knockout decreases SMAD1 phosphorylation and Id1 expression while *FYN* knockout promotes BMP9 mediated Id1 expression and increases SMAD1 phosphorylation under certain media conditions (Sup. Figure 2E,F). It is noted the effect was studied in several media conditions. Changes in mRNA could be demonstrated with cells cultured in full endothelial growth media (EGM). Changes in protein expression required starvation in 10% EGM prior to stimulation.

**Figure 3:**
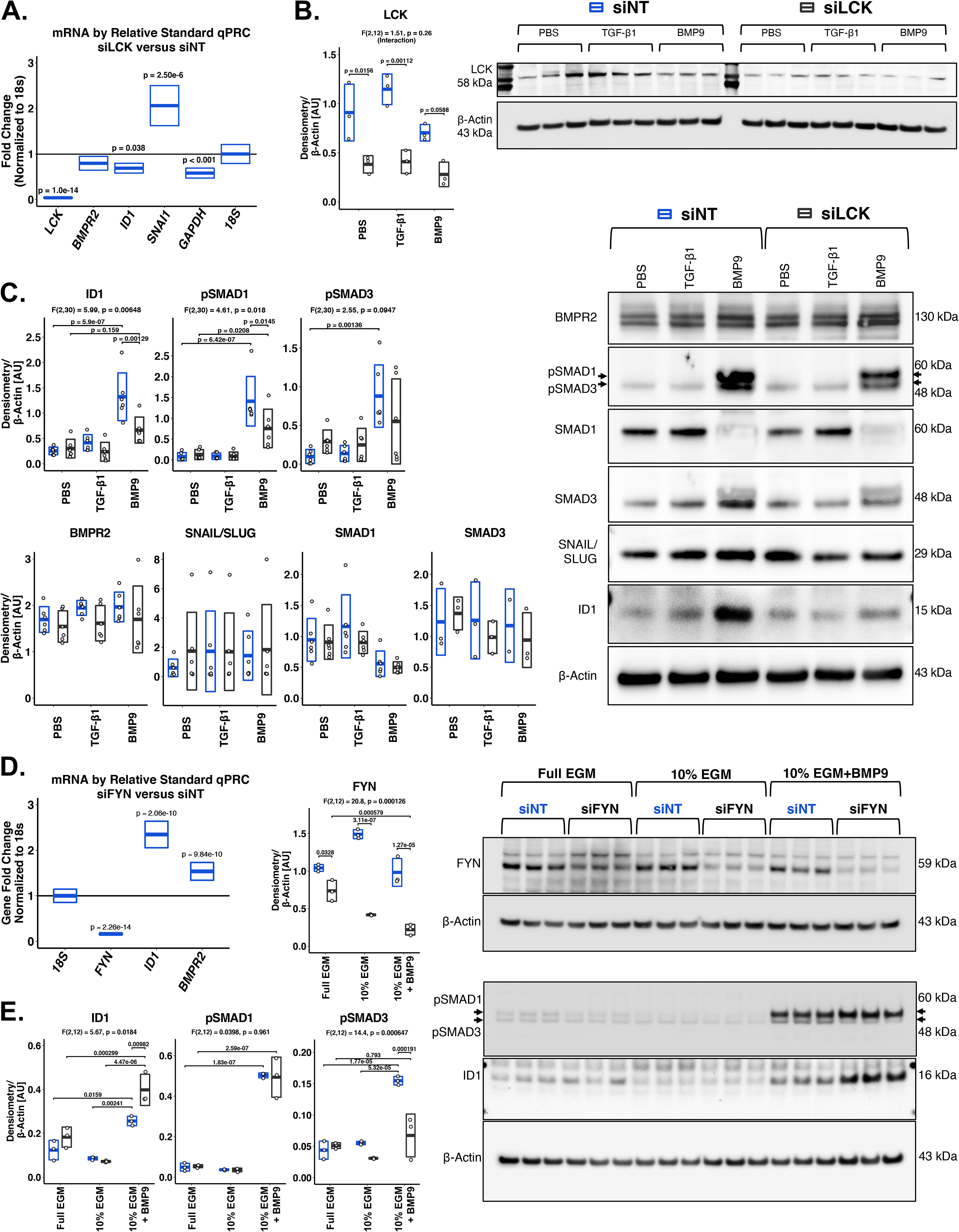
Silencing of the SrcB LCK in human PAECs suppresses canonical BMPR2 signaling, whereas silencing the SrcA FYN supports it. (A) Human PAECs transfected with *LCK* targeting (siLCK) or control (siNT) siRNA and lysed at 72 hours in full endothelial growth media (EGM) have decreased *LCK*, *ID1*, and *GAPDH* by RT-qPCR using the relative standard method. *SNAI1* was significantly increased. 1-Way ANOVA with Tukey’s Rage Test, n = 4. **(B)** siLCK PAECs have lower LCK expression at 72h in full EGM. **(C)** SiLCK PAECs have significantly decreased BMP9 mediated phosopho-SMAD1 and ID1 expression. There was no change in BMPR2 or total SMAD1 expression across all siLCK conditions. N = 6, 2-way ANOVA for treatment (PBS, TGF- β1 and BMP) and transfection (siNT vs. siLCK) by Tukey’s Range Test. **(D)** PAECs transfected with siFYN or siNT for 6 hours, starved at 24 hours in 10% EGM, harvested at 72 hours, have significantly decreased FYN, but increased ID1 and BMPR2 using RT-qPCR by relative standard method with normalization to 18s RNA. Analysis by 1-way ANOVA, n = 4. **(E)** siFYN PAECs maintained in full EGM, changed to 10% EGM 24 hours after transfection, or changed to 10% EGM then given BMP9 or PBS 1.5 hours prior to cell lysis. FYN was significantly reduced at 72 hours. There was no change in Phospho-SMAD1 activity (pSMAD1). There was significant reduction in phospho-SMAD3 in siFYN, BMP9-treated cells and a significant increase in Id1. 2-way ANOVA examining treatment (Full EGM, 10% EGM + PBS, 10% EGM+BMP9) and transfection (si-NT vs. si-FYN) followed by Tukey’s Range Test. For all above ANOVA tests, normality checks for ANOVA residuals and Levene’s test for homogeneity were carried out and assumptions are met.

### *LCK* but not *FYN* suppression is associated with EC dysfunction in human PAECs

Next, we asked if the changes in BMPR2 signaling due to *LCK* or *FYN* knockout associate with phenotypic changes in PAECs. SiLCK and siFYN PAECs were examined in tube formation angiogenesis assays alongside non-targeted siRNA treated (siNT) PAECs. Because the optimal concentrations of siRNA and lipofectamine required to achieve an effective knockdown of *LCK* in PAECs is different than the optimal concentrations for *FYN*, siLCK and siFYN cells are compared to siNT controls termed siNT-F and siNT-L that have matched siRNA and lipofectamine concentrations. Viable cells were seeded at a constant density on Matrigel 48-hours post transfection. Cells undergoing *FYN* knockout had increased tube formation with a higher total tube length and more tube junctions relative to siNT-F (Figure 4A-B). Conversely, siLCK PAECs sprouted few, shorter tubes and decreased tube junctions compared to siNT-L (Figure 4A-B). Notably, cell death was accelerated in siLCK versus siNT-L cells (Figure 4C). We hypothesized this correlated to an increase in PAEC apoptosis. Indeed, we found higher *BAX* and *BCL* levels in siLCK versus siFYN cells relative to their respective controls (Figure 4D). Increased apoptosis was confirmed by assessing Caspase 3/7 activity (Figure 4E). Because EC dysfunction includes upregulation of integrins along with other pro-inflammatory cues, we assessed VCAM1 expression 72 hours after siLCK and siFYN transfection in full EGM, finding an increased expression of VCAM1 in siLCK versus siFYN cells (Figure 4F). Taken together, this suggests that suppression of the SrcB LCK generates a dysfunctional EC phenotype as opposed to the SrcA FYN.

**Figure 4:**
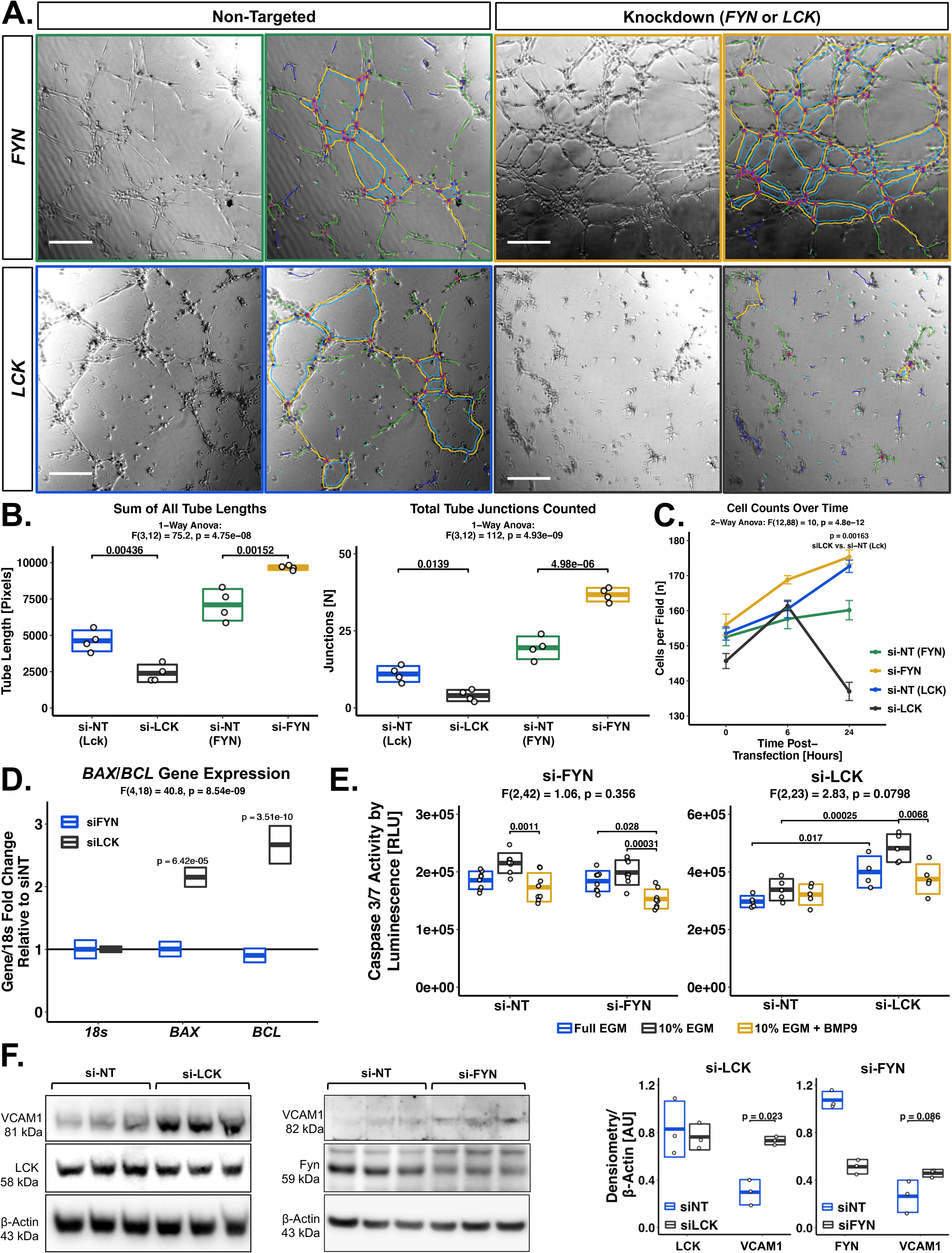
Inhibition of the SrcB LCK promotes EC dysfunction, whereas inhibition of the SrcA FYN does not. siLCK and siFYN PAECs are compared to matching control conditions (termed si-NT-L or si-NT- F). Cells were passaged to Matrigel in full endothelial growth medial (EGM) 48 hours after transfection and imaged 3 hours later. **(A)** Raw (left) and analyzed (right) brightfield images are shown. **(B)** Total tube length and junctions are decreased in siLCK PAECs. Conversely, siFYN PAECs have increased total tube length and junctions. N = 4 experiments. **(C)** siLCK PAEC cell counts in culture decrease over time. **(D)** RT-qPCR by relative standard methods in siLCK and siFYN PAECs in starvation media (10% EGM) at 72 hours shows a significant increase for BAX and BCL mRNA relative to 18s mRNA. 1-Way ANOVA with Tukey’s range test. N = 4. **(E)** Increased (siLCK vs. siNT-L) PAECs and decreased (siFYN vs. siNT-F) Caspase 3/7 activity at 48 hours post transfection in 10% EGM. **(F)** siLCK PAECs had increased expression of VCAM1 despite minimal reduction in LCK protein 72 hours after transfection in full 10% EGM media. Statistical comparison with Student’s T-Test.

### Whole transcriptome analysis finds *LCK* and *FYN* co-regulate genes that group into two distinct signatures. One signature supports endothelial homeostasis and includes Bmpr2 pathway members. The second signature is inflammatory and includes canonical NF-κB responsive genes

To this point, we have taken a focused view examining canonical BMPR2 signaling in response to *LCK* and *FYN* knockout in PAECs. As each PTK engages thousands of tyrosine phospho-targets in eukaryotic cells, it is unlikely that LCK and FYN influence endothelial function only through modulating of BMPR2 signaling. To determine if additional signaling pathways outside of the canonical BMPR2 signaling pathway are differentially regulated by LCK and FYN we performed a transcriptome-wide assessment and identified two sets of genes differentially regulated by LCK and FYN which we term “endothelial” and “inflammatory” signatures.

Bulk-cell RNA sequencing was conducted in four groups of PAECs: siLCK, siFYN, a scrambled siRNA control group (NT), and a scrambled siRNA group subjected to BMP9 stimulation (siNT + 50 ng/mL BMP9 for 2 hours prior to harvest). For all arms, the same concentration of siRNA and lipofectamine was used to control for gene expression variability induced by the transfection process. Gene knockdown was confirmed by RT-qPCR prior to sequencing. Principal component analysis demonstrates clustering of replicates with between group separation (Sup. Figure 3B). Volcano plots of differentially expressed genes (DEGs) for siLCK and siFYN conditions relative to siNT are shown in Sup. Figure 3C,D. *FYN* was identified as a significantly downregulated gene in the siFYN arm. Although *LCK* was not found to be statistically downregulated in the RNA seq siLCK arm due to low baseline counts (10-15 raw reads), its knockdown was confirmed by RT-qPCR.

From the heatmap in Figure 5A, we identified *Id1, Bmpr2,* and other Bmp-responsive genes (*SMAD6*, *Hey2*) [31] in a group increased by *FYN* knockout and decreased by *LCK* knockout. Additionally, a large set of genes were increased by *LCK* knockout and decreased by *FYN* knockout and implicated an inflammatory and dysfunctional endothelial phenotype: *Il1B* (IL-1β), *VCAM1*, *CCL8*, and *CXCL3* [32]. We noted that genes associated with endothelial identity (*Pecam1*, *Gja5*, *Lyve1, Cdh5*, *Cldn5*) [33] were simultaneously decreased by *LCK* knockout and increased by *FYN* knockout (Figure 5C and Sup. Figure 4).

**Figure 5:**
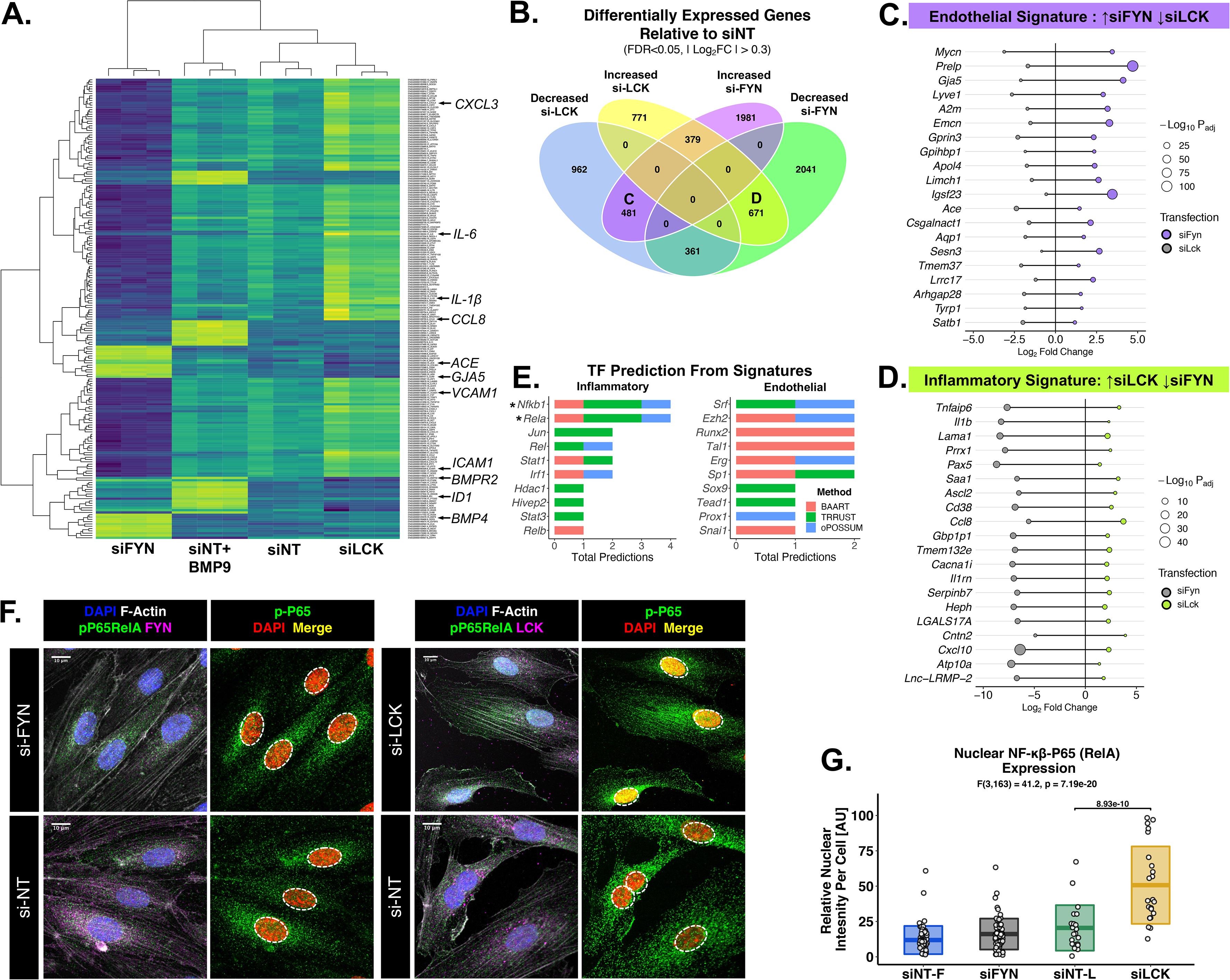
Whole-transcriptome analysis finds LCK knockout suppresses, and FYN knockout promotes an “endothelial” gene signature (including the BMPR2 pathway). Conversely, LCK knockout increases and FYN knockout suppresses an “inflammatory” gene program centered on NF-κB / RelA signaling. (A) Normalization of raw counts by variance stabilizing transformation with unsupervised hierarchical clustering and heatmap generation. Genes increased by *FYN* knockout and increased in *LCK* knockout correlate with canonical BMPR2 signaling (*BMPR2*, *ID1*, *BMP4*) and endothelial identity (*GJA5*, *ACE*). Genes increased by *LCK* knockout and decreased by *FYN* knockout include inflammatory and ECD related genes (*CXCL3*, *IL-6*, *CCL8*, *VCAM1*, and *ICAM1*). (B) DEGs co-regulated by *FYN* and *LCK*. 671 genes increase with *LCK* knockout and decrease with *FYN* knockout (Group D, “Inflammatory” signature). 481 genes decrease with *LCK* knockout and increase with *FYN* knockout (Group C, “Endothelial” signature.) (E) Transcription factor (TF) analysis shows inflammatory signature DEGs center on canonical NF-κB/p65 RelA signaling whereas multiple TFs are implicated by the endothelial signature. (F,G) IF staining of siLCK, siNT-L, siFYN, and siNT-F PAECs verifies increased pp65 RelA nuclear co- localization in siLCK PAECs. Statistical analysis with 1-Way ANOVA.

Because *LCK* and *FYN* knockout exert differential control over endothelial phenotypes, we decided to focus on the set of genes that are co-differentially regulated by *LCK* and *FYN.* The Venn diagram in Figure 5B finds that 481 genes were decreased by *LCK* knockout *and* increased by *FYN* knockout, whereas 671 genes were increased by *LCK* knockout *and* decreased by *FYN* knockout. Gene set enrichment analysis (GESA) [34] and transcription factor prediction [35–37] was performed of both these signature (Sup. Figure 5).

The 671 genes increased by *LCK* and decreased by *FYN* were found by GSEA to associate with a type 1 interferon or lipopolysaccharide response. Further, there is enrichment of genes associated with interferon α/β stimulation, NF-κB activation, and MHC-1 activation (Sup. Figure 5D,E). Indeed, transcription factor prediction by three methods found that the canonical NF-κB transcription factor *RelA* linked a significant proportion of genes in this group (Figure 5E). This finding was confirmed in cell culture, where phosphorylated p65-RelA was found to be significantly increased in the nuclei of PAECs subjected to *LCK* knockout, but not *FYN* knockout (Figure 5F, 5G). Given this, we refer to this set of 671 genes as an “inflammatory signature” linked by canonical NF-κB activity.

For the 481 genes increased by *FYN* knockout and decreased by *LCK* knockout, GSEA analysis found significant enrichment of genes involved in sprouting angiogenesis and endothelial cell migration (Sup. Figure 5B,C). The overall signature was more complex, with transcription factor analysis implicating *Sp1, Erg, Runx2,* and *Snai1* (Figure 5E). *Erg* is decreased in PAH [38] and exerts control over Notch signaling to promote vascular stability [39]. *Sp1* activates VEGF expression in tumor derived ECs and acts as a promoter for the gene *Endoglin. Sp1* loss can contribute to HHT and PAH via *Endoglin* suppression [40]. Based on the GSEA results, the aforementioned decrease in endothelial-specific genes (e.g. *Gja5*, *Ace*), and the demonstrated suppression in Bmpr2 signaling, we term this set of genes an “endothelial signature” associated with EC identity, BMPR2 signaling, and endothelial migration

In summary, whole transcriptome analysis after loss of function of *LCK* (Src-B) and *FYN* (Src-A) in PAECs finds a co-regulated set of genes that can broadly be divided into an “endothelial” signature (including Bmpr2 signaling) and an “inflammatory” signature (including NF-κB signaling).

### Querying the “Endothelial” and “Inflammatory” gene signatures in the CLUE Connectivity Map (CMap) database finds TKIs that mirror the effects of selective SrcA and SrcB knockout

To this point we have manipulated *Lck* and *Fyn* through transfection of siRNA to illustrate the different functions of SrcA and SrcB kinases in PAECs. We next wanted to find tyrosine kinase inhibitors (TKIs) that could exploit this functionality in endothelial cells. However, a major barrier is the fact that TKIs are promiscuous with respect to PTKs and seldom target a single kinase. Because the number of kinase substrates exceeds PTKs by a factor of 100, a nonspecific PTK-TKI interaction has the potential to unpredictably influence multiple cellular signaling pathways. To circumvent this, we hypothesized that using transcriptomic data representing the cell states described by the endothelial and inflammatory signatures could help predict drugs that mimic the effect of selective SrcB or SrcA knockout. The availability of Broad Institute CLUE Connectivity Map (CMap) database and search tools (a superset of gene signatures which includes NIH LINCS/L1000 data) allows us to link perturbagens (genes manipulations, ligands treatments, and small molecules) to our endothelial and inflammatory signatures [41]. The search strategy and resulting perturbagens are shown in Figure 6A.

**Figure 6:**
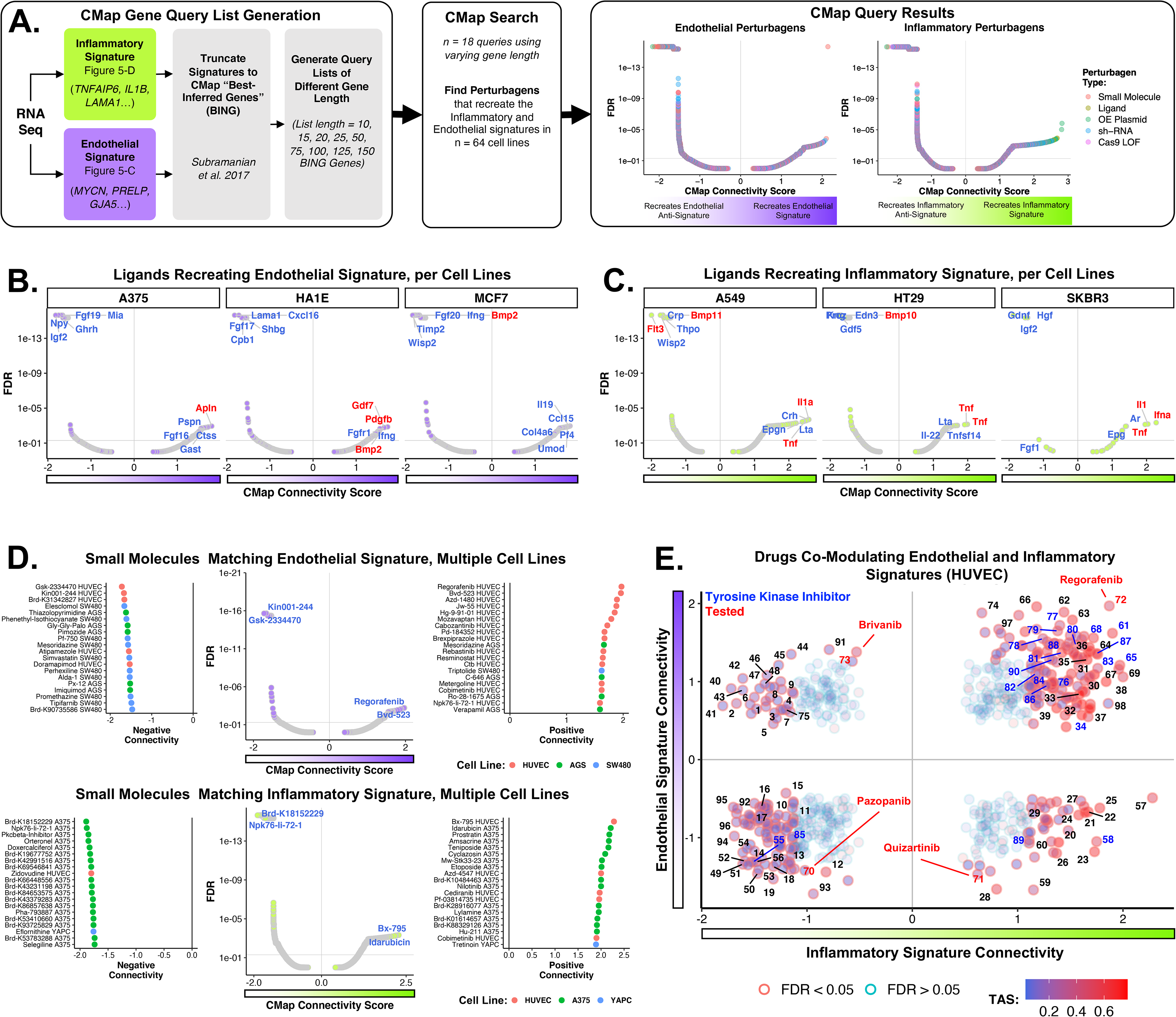
Using endothelial and inflammatory signatures, CMap searches validate ligands and enables drug prediction. (A) Search strategy for perturbagens in the CMap data with all perturbagen results. **(B)** Ligands that generate high connectivity scores with the endothelial signature in select cell lines. Note BMP2 can have high positive or negative connectivity depending on the cell line. **(C)** Ligands generating high connectivity scores with the inflammatory signature. Note that TNF has high positive connectivity with the inflammatory signature across cell lines. **(B,C)** Top five ligands with the highest connectivity scores are annotated, red indicates a ligand of interest. **(D)** Top compounds with high connectivity to endothelial (top) and inflammatory (bottom) signature/anti-signature in cell lines. **(E)** Compounds which generate high connectivity scores for both the endothelial and inflammatory signatures in HUVEC cells. Blue text denotes TKIs which are most prominent in the upper right inflammatory and endothelial quadrant. Red indicates are representative of the four “quadrants”. FDR = false discovery rate. TAS = transcriptional activation score, a measure of signature strength and replicate correlation intended to represent a perturbagen’s transcriptional activity.

There are several barriers to using CMap. First, our gene signatures needed to be trimmed to include genes that are measured or accurately inferred (termed “Best INFerred” or BING) on the L1000 platform [41]. Second, the number of genes to use in the CMap search query (ranging from 10 to 150) required optimization. Third, it was unclear which of the immortalized or neoplastic CMap cell lines would be the best substrate for human endothelial cell drug prediction. We detail the handling of list generation and gene list length selection in the methods and Sup. Figure 6A.

As shown in Figure 6A, CMap returns perturbagens that “connect” to our queried endothelial and inflammatory signatures. Perturbagens with a high positive connectivity score induces the signature (upregulation of queried signature genes) whereas negative connectivity score induces the anti-signature (downregulation of queried signature genes) in CMap cell lines. We first studied the ligand perturbagens to see if ligands with high connectivity aligned with our prior experiments. In Figure 6B, the ligands BMP2, GDF7 (a TGF-β ligand), and Apelin had strong positive connectivity with the endothelial signature in A375 (melanoma) and HA1E (kidney) cells. However, BMP2 also had strong negative connectivity in MCF7 (breast cancer) cells. We drew two conclusions from this. First, CMap independently validates that the endothelial signature is a Bmp-responsive signature. BMP2 in HA1E CMap cells upregulates the same genes as *FYN* / SrcA knockout did in PAECs. Second, the endothelial signature is plastic across cell lines – the same ligand (BMP2) can induce or suppress the same gene signature depending on the cell line being treated (Results for all CMap cell lines are shown in Sup. Figure 6G).

This contrasts to the inflammatory signature in Figure 6C. Here, strong positive connectivity was seen between the inflammatory signature and the ligand Tumor necrosis factor (TNF) in a majority of cell lines (All cell lines shown in Sup. Figure 6F). Because TNF is an inducer of NF-κB [42] these results align with our prior finding that LCK / SrcB knockout activates NF-κB. Importantly, conserved TNF connectivity with the inflammatory signature contrasts with the dichotomous and plastic connectivity between Bmp2 and the endothelial signature.

Given the above heterogeneity, we took two approaches to determine which cells would be the most suitable cell substrate for drug prediction. The first was a pragmatic approach where we selected cell lines that gave the highest positive connectivity scores with the endothelial signature and were biologically plausible. This quickly led us to the HUVEC line, an endothelial cell line which in Figure 6D had the strongest endothelial signature connectivity of any cell line for small molecules. For the second method, we leveraged the fact that some CMap cells underwent *LCK* and *FYN* knockout and overexpression experiments. We selected those CMap cells that had strong positive connectivity to PAECs subjected to *LCK* and *FYN* knockout in our previous experiments (Sup. Figure 6B,C). Interestingly, we found two populations of cells: ones that had strong positive connectivity to *LCK* or *FYN* knockout in PAECs (HEK293T kidney and AGS gastric cells) and ones that had strong negative connectivity (HCC515 adenocarcinoma for *FYN* and VCAP prostate cancer for *LCK*). We concluded that knockout of the same gene can produce either a signature or anti-signature depending on cell substrate, further underscoring the importance of cell line choice in drug prediction.

Turning to small molecule perturbagens, we found that some agents, such as the TKI Regorafenib, connected strongly with both the endothelial signature and inflammatory signatures. We hypothesized that if we tested these compounds intending them to increase genes in the endothelial signature that they may also have an unanticipated inflammatory affect. Therefore, we decided to investigate small molecule perturbagens considering their connectivity to both the endothelial and inflammatory gene signatures simultaneously. We did this analysis in the HUVEC line (due to the number of perturbagens with high connectivity scores) and focused on TKIs (Figure 6E). Notably, a majority of TKIs had positive connectivity with both the inflammatory and endothelial signatures (top right corner, Figure 6E). Few TKIs existed in what we presumed to be a more desirable quadrant – positive endothelial and negative inflammatory connectivity (top left corner, Figure 6E). We set out to determine how representative compounds from each quadrant affected PAECs (all compounds annotated in Sup. Figure 7).

**Figure 7:**
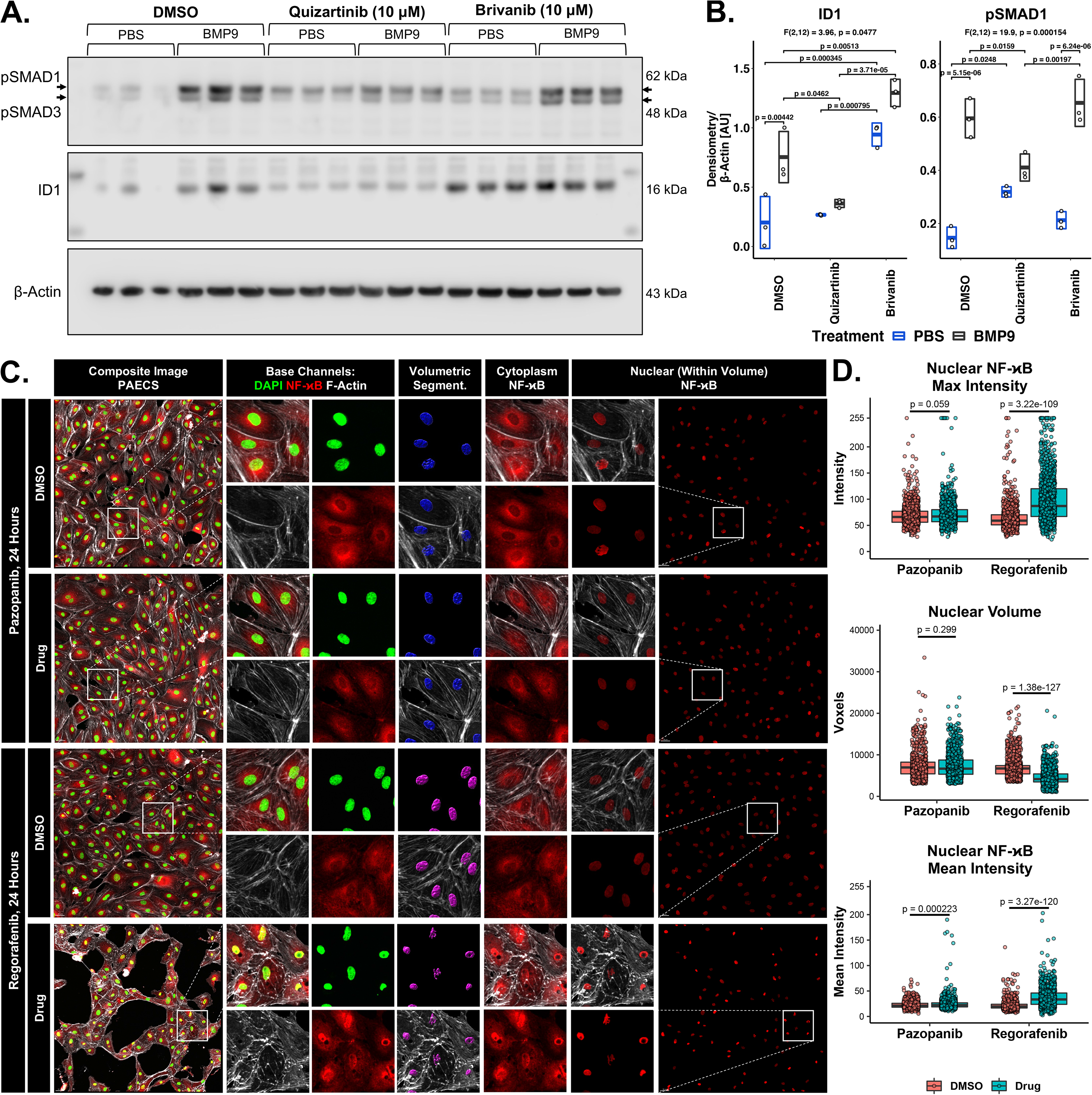
Brivanib and Quizartinib increase and decrease PAEC ID1 expression, whereas Regorafanib increases PAEC NF-. κ**B activity**. **(A)** PAECs in full EGM treated with drug or DMSO for 24 hours then given BMP9 or PBS 2 hours prior to harvest significantly increase ID1 expression with Brivanib alone or Brivanib plus BMP9 versus DMSO. Quizartinib decreases ID1 in all cases. Statistics by 2-way ANOVA investigating the interaction between drug and BMP9 treatment with Tukey’s Range Test, n = 3. **(B)** PAECs treated with Regorafenib, Pazopanib, or DMSO for 24 hours in full EGM were fixed and stained for p65 RelA, F-Actin, and DAPI. DMSO controls were paired on the same slide to control for irregularities in the staining. Cell barrier integrity was compromised in Regorafanib treated cells. **(C)** The average (average of p65 intensity across each voxel contained in the nucleus) and maximum nuclear p65-RelA expression was increased in Regorafenib treated PAECs relative to DMSO. Nuclear size was significantly decreased in Regorafenib treated cells relative to DMSO treated cells. Statistics by Student’s two-tailed T-test.

Ten compounds underwent a preliminary screen in PAECs. The most promising candidates from preliminary screens were selected from each quadrant: Regorafenib (top right), Brivanib (top left), Pazopanib (bottom left) and Quizartinib (bottom right). In Figure 7A-B, we examine if Quizartinib and Brivanib generate the anticipated endothelial signature effect by examining SMAD phosphorylation and ID1 expression in PAECs, with or without BMP9 stimulation. Pre-treatment with Quizartinib in DMSO versus DMSO alone for 24 hours fully suppressed expression and significantly decreased p-SMAD1 and ID1 expression 2 hours after treatment with BMP9. Conversely, Brivanib increased basal expression of ID1 as well as BMP9 mediated expression of ID1 but did not have a significant effect on p-SMAD1 expression. The augmentation of ID1 without pSMAD1 increase was similarly observed with *FYN* knockout, Figure 3E. We therefore show that two drugs predicted by our search method, Quizartanib and Brivanib, exhibit the anticipated effect on ID1.

In Figure 7C, we examine how Regorafenib and Pazopanib generate the anticipated inflammatory signature using NF-κB signaling modulation as a read-out. Regorafenib treatment resulted in disruption of the EC barrier with increased nuclear p65 RelA expression (Figure 7D) and decreased nuclear size. Thus, Regorafanib increases canonical NK-κB signaling as predicted by our CMap searches. Pazopanib had little effect on NF-κB signaling or the EC barrier, possibly due to low baseline activity or lack of direct NF-κB stimulation. It is notable that despite the predicted endothelial effect of Regorafanib, the inflammatory signature appears to “win out”, negating the endothelial effect.

## Discussion

This study is significant for several reasons. The key finding is that Src-A and Src-B protein tyrosine kinases differ not only by phylogenetic classification, but also in how they modulate cellular signaling pathways. When granularly focusing on the example SrcA and SrcB kinases FYN and LCK, we find differential regulation of both the BMPR2 and the NF-κB pathways occurs in PAECs. Second, although prior studies have investigated the effect of Src-Family PTKs in the pulmonary vasculature, we specifically investigate FYN and LCK in PAECs, a lineage implicated at the inception of lung diseases such as PAH. By showing selective SrcB (LCK) PTK knockout both blocks BMPR2 and increases NF-κB signaling we provide an additional rational to link TKIs to the development of EC dysfunction through inhibition of SrcB kinases. Third, we use CMap data to predict drugs mimicking the effects of FYN and LCK inhibition in PAECs. We reach several conclusions about the drug prediction methodology. First, a transcriptomic signature can be both upregulated or downregulated by the same perturbagen in different CMap cell lines. This argues against indiscriminately using all available cell lines for drug prediction. Instead, by tailoring the search to include biologically congruent cell lines, we recover ligands and drugs matching our predicted outcome. Second, we find that the same drug can simultaneously affect two gene signatures, with one signature (e.g. inflammatory) “wining out” over another (e.g. endothelial) in testing. Thus, it may be more fruitful to perform transcriptomic drug prediction using several desired gene signatures (e.g. pro-endothelial and anti-inflammatory).

### SrcA and SrcB Kinases Differentially Regulate Cell Signaling Pathways

Because 518 kinases phosphorylate an estimated 119,809 unique substrates [43], PTKs must be regulated to avoid unintended signal activation via crosstalk. For Src kinases, it was believed this was achieved via cell- specific expression, with SrcA expressed in diverse cell types and SrcB restricted to hematopoietic lineages [44]. Phylogenetic analyses now show that SrcA and SrcB are genetically distinct [1]. Further, the amino acid sequence of the kinase pocket also obeys the SrcA and SrcB phylogenic division and confers kinase-substrate specificity [45]. This is believed to prevent kinase crosstalk in regions where PTKs are in close spatial proximity, as with LCK and FYN in the T-Cell Receptor [46, 47]. To our knowledge, we are the first to demonstrate that cellular signaling pathways react in differently to selective SrcA and SrcB knockout. Thus, SrcA and SrcB PTKs can be distinguished by genetic sequence, kinase pocket amino acid sequence, and in how they support or inhibit cellular signaling pathways. It is highlighted that our agnostic siRNA screen found all SrcA and SrcB kinases modulate canonical BMPR2 signaling in the same way, albeit at different levels.

We narrow to two example PTKs and show this effect holds in human PAECs and extends to the NF-κB pathway. It is likely that other BMP signaling pathways comprised of different type 1 and type 2 TGF-β receptors are differentially regulated by SrcA and SrcB family members. A next step could be to examine the effect of LCK and FYN on BMP signaling in lymphocytes using *Id2* as the BMPR2-responsive readout, based on our public sc-RNA seq PBMC analysis. Finally, the exact mechanism for FYN and LCK regulation of BMPR2 and NF-κB are not fully explored. Hypotheses include FYN and LCK kinases potentiating conformational changes in SMAD proteins or their inhibitors [30], FYN and LCK acting as a scaffolding proteins for membrane receptors, or FYN and LCK modulating the expression of BMP ligands within culture (due to our observation of increased *BMP4*, *BMP12* and *BMP13* mRNA by RNA seq).

### LCK and FYN are present in PAECs. Selective inhibition of the SrcB LCK can drive EC dysfunction

Studies have investigated the role of Src kinases in the pulmonary vasculature, emphasizing smooth muscle cells (SMCs) [26, 48]. Notably, LCK is not found at high levels in SMCs and few studies have investigated LCK in ECs [49]. Here, we show LCK is expressed at a message and protein level in PAECs and knockout of *LCK* has a functional consequence in human PAECs. Notably, we were unable to detect LCK protein or mRNA in the ECs of mice or rats, by immunostaining or from public RNA seq data sets. LCK and FYN are present in lymphocytes and LCK levels are reduced in PBMCs of patients with interstitial lung disease [50], lupus [51], and, as we have shown, pulmonary arterial hypertension [24]. However, as shown in Figure 2, the more ubiquitous peripheral reduction of LCK in PAH patients does not translate to the lung endothelium in PAH patients. We did however observe a correlation between LCK and Id1 expression in cultured patient PAECs, implying LCK may positively facilitate BMP signaling in ECs. As LYN is expressed in SMCs, future studies could use LYN as representative SrcB kinases, potentially comparing to the known effect of SRC [26, 52].

### Using endothelial and inflammatory gene signatures, we employ a biologically tailored methodology of drug prediction to manipulate canonical BMPR2 and NF-κB signaling in PAECs

Drugs are inherently dirty and can modulate several signaling pathways simultaneously. TKI-PTK kinome analyses shows that despite heterogeneity, conservation of the ATP-kinase-pockets of PTKs results in promiscuous PTK inhibition by TKIs and few TKIs that are truly specific for a single PTK target [53]. The kinase pockets of SrcA and B kinases are distinct [45] but still share homology. It is therefore a reasonable assumption that a TKI will bind many PTKs simultaneously in a cell with varying affinity, influencing several pathways at once. To manage this, we hypothesized that instead of using protein-TKI interactions we could instead find TKIs by focusing on a desired transcriptomic cell state. By using our FYN- and LCK-derived endothelial and inflammatory signatures, we aimed to find TKIs with inherent affinities for SrcA or SrcB PTKs.

We leveraged the Broad institute’s CLUE CMap database, noting that methodology around its usage is evolving [54]. We make several observations. First, CMap query output was dependent on the number of BING genes used in the search. A unique number of search genes retrieves a maximal number of perturbagens across all CMap cell lines – on the order of 5x10^5^. We found the endothelial and inflammatory gene signatures derived from PAECs could be both induced or suppressed by the same ligand or gene manipulation depending on the CMap cell line the perturbagen was used in. Many cell lines in CMap (and LINCS) are immortalized cancer lines whose genetic programs may differ from healthy cell lines. To date, studies using LINCS to predict drugs from, for example, PAH or Ulcerative Colitis gene signatures have not restricted by cell line [55, 56]. Our data suggests cell line choice is important. A biologically tailored approach to match the substrate cell line with the experimental cell line may focus the results.

Finally, it should be said that Brivanib, Quizartinib, and Regorafanib are used as “proof of concept” TKIs to validate our search strategy. Brivanib interestingly targets VEGFR1,2 and FGFR1 to inhibit angiogenesis in cancer models [57]. Quizartinib is a selective FLT3 inhibitor that targets PDGFR, and VEGFR2 to a lesser degree [58]. Regorafanib is a multi-kinase inhibitor targeting VEGFR1,2, KIT, BRAF, and TIE1 among others [59]. Further pre-clinical work is needed to determine if the observed effects on canonical BMPR2 expression in quiescent large vessel PAECs by Brivanib has a role in the treatment of human pulmonary vascular diseases.

## Methods

### 1. High Throughput siRNA Screen

High throughput siRNA screening of > 22,000 genes using an Id1-BRE luciferase reporter assay in a C2C12 mouse myoblastoma cell line treated with or without 250 pM BMP4 was conducted in the Stanford High-Throughput Bioscience Center, as previously described [60]. Briefly, the C2C12 myoblastoma cells, a generous gift from Dr. Peter ten Dijke, were stably transfected with BRE- Id1 linked to luciferase and used as a reporter cell line [25]. Cells were screened on 384-well siRNA plates. The transfection conditions were optimized with BMP4 as stimulus, siBMPR2 and a toxic (siTox) control. Cells were seeded in wells at a density of 1500 cells/well. Next, siRNA was transfected at an optimized concentration of 25 nM using DharmaFect3. After 48 hours, cells were stimulated with 250 pM of BMP4. Two hours after stimulation, the change in luminescence of an individual well was measured relative to the average luminescence of all wells in the plate. After this, a tryptan-blue cell viability stain was automatically performed and quantified. Luminescence and viability stain intensity was normalized to the overall average luminescence and viability stain intensity of the plate to calculate change in *Id1* expression and change in viability. This assumes that there are few “hits” (e.g. few significant deviations in *Id1* expression or cell viability from the norm) per 384 well plate and is a standard method for batch effect normalization in multi-plate high throughput assays. Transfections were performed in triplicate for each gene. Change in Id1 linked luminescence relative to baseline and change in cell viability relative to baseline was averaged for three samples across all 22,124 genes used in the high throughput murine wide si-RNA screen.

### 2. Cell Culture

Human large vessel PAEC (Promocell) were grown as monolayers in 0.2% gelatin in PBS coated dishes in a commercial Promocell Endothelial Cell Growth Media (Cat No. C-39210). Contents of full Endothelila Growth Media (EGM) are: fetal calf serum (0.02 ml/ml), recombinant human epidermal growth factor (0.1 ng / ml), recombinant human basic fibroblast growth factor (1 ng/ml), heparin (90 μg/ml), hydrocortisone (1 μg/ml), and bovine hypothalamus extract (0.004 ml/ml). Cells were passaged at 1:3 ratios and used for experiments at passages of between 3-6. For starvation, full EGM was diluted by 10% in basal growth media (10% EGM).

### 3. Human PAH Tissue: Isolation of Cells lung tissue

PAEC of IPAH and FPAH patients at time of lung transplant were obtained from digested whole lung tissue, using CD31-AB pulldown beads (Dynabeads; Invitrogen), as previously described [60]. Experiments involving human tissue or derived primary cells were approved by the Stanford University Institutional Review Board and the Administrative Panel on Human Subject Research.

### 4. RNA Interference

Both *Lck* and *Fyn* expression was modulated by RNA interference in PAEC. A pool of 4 siRNAs for BMPR2, FHIT, LCK or a non-targeting control pool (Ambion, Table. 1) were transfected into PAECs using the RNAi Max kit (Invitrogen) for 48 hours. mRNA knockdown efficiency was determined by qPCR.

All transfections were carried out in 6 well plate format. Transfection conditions were varied to independently optimize transfection efficiencies for *Lck* and *Fyn*, resulting in different concentrations of siRNA and Lipofectamine for each gene. First, cell media was changed to 500 μL of OptiMem Media (Thermo Fisher 31985062) 30 minutes prior to transfection. For si-Lck conditions, 4 μL of Lipofectamine (RNA iMax, Invitrogen 13778) was added to 246 μL of OptiMem and 4 μL of 20 nM siRNA (either *Lck* or nontargeting, NT) was added to 246 μL of OptiMem. The Lipofectamine and siRNA containing OptiMem was combined. After letting stand for 20 minutes, this 500 μL was added existing OptiMem in 6 well plates. This resulted in a final concentration siRNA concentration of 80 nM. For si-Fyn, the procedure was the same except 2 μL of Lipofectamine in 248 μL OptiMem and 2.5 μL of siRNA (either *Fyn* or NT) in 247.5 μL OptiMem was used to make a final siRNA concentration of 50 nM. All analyses for each gene were done by comparing to the same concentration of NT siRNA. For the RNAseq experiments, to control for transfection mediated gene effects, the same conditions used for siFYN above were used for the transfection of *Fyn*, *Lck*, and NT siRNA. 50 nM of siRNA and 2 μL Lipofectamine in 1000 μL OptiMem was used for all experimental arms.

### 5. qPCR Assay to Detect mRNA Expression

For mRNA, total RNA was extracted from adherent cell layers using the RNAeasy Plus Kit (Qiagen) and reverse transcribed into cDNA using random primers with the Taqman cDNA reverse transcription Kit (Applied Biosciences) according to the manufacturer’s instructions. Prior to the generation of cDNA, RNA quality and concentration was measured using the Nanodrop system (Thermo-Fisher). The level of mRNA expression was quantified using Taqman primer/probe sets (Table 2) for the target and normalized to a housekeeping control (18s). To validate new PCR probes, standard curves were calculated from serial 1:2 dilutions of cDNA across 6 different concentrations, and PCR probe efficiency (E) was calculated. From the standard curve, the relative standard method was used to determine relative concentrations of mRNA in lysates. Note that change in *Gapdh*, *18s,* and *B2M* genes were examine relative to one other. For *LCK* conditions, *Gapdh* was found to be significantly downregulated relative to be both *18s* and *B2M* genes (results for *18s* shown in Figure 3A). Thus *18s* was selected as the housekeeping gene for RT-qPCR experiments. The change in *Gapdh* was not observed with *FYN* knockout.

**Table 1:**
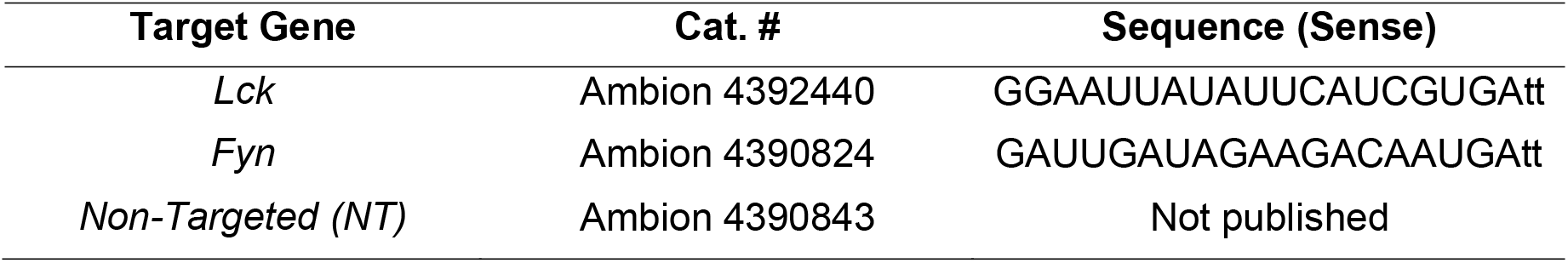

**Table 2:**
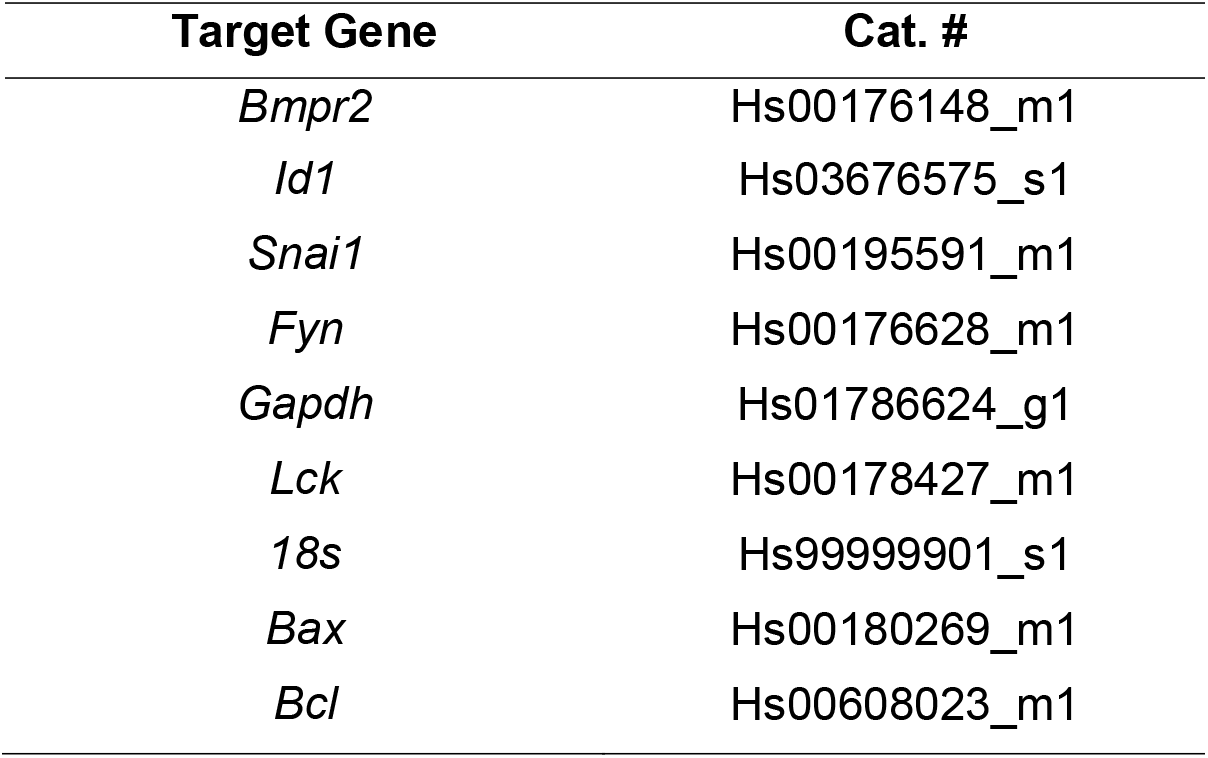

| Target Gene | Cat. # |
| --- | --- |
| *Bmpr2* | Hs00176148_m1 |
| *Id1* | Hs03676575_s1 |
| *Snai1* | Hs00195591_m1 |
| *Fyn* | Hs00176628_m1 |
| *Gapdh* | Hs01786624_g1 |
| *Lck* | Hs00178427_m1 |
| *18s* | Hs99999901_s1 |
| *Bax* | Hs00180269_m1 |
| *Bcl* | Hs00608023_m1 |

### 6. Western Blotting

Western blotting was performed as previously described [60]. Antibodies were used as described in Table 3:

**Table 3:**

| Target | Manufacturer | Cat. # | Clone | Species | Dilution |
| --- | --- | --- | --- | --- | --- |
| β-Actin | Santa Cruz | SC4778 | C4 | Mouse | 1:600 |
| BMPR2 | Invitrogen | MA5-15827 | 3F6F8 | Mouse | 1:800 |
| Lck | Abcam | Ab227975 | EPR20798-107 | Rabbit | 1:300 |
| Id1 | Santa Cruz | sc-133104 | B8 | Mouse | 1:100 |
| Snail/Slug | Abcam | Ab180714 | Polyclonal | Rabbit | 1:500 |
| VCAM1 | R&D Systems | BB16-V1 | BBIG-V1 | Mouse | 1:500 |
| Fyn | BD Biosciences | BD61063 | 25/Fyn | Mouse | 1:500 |
| pSMAD-1/3 | Abcam | ab52903 | EP823Y | Rabbit | 1:1000 |
| SMAD1 | Cell Signaling | CST 9743 | Polyclonal | Rabbit | 1:500 |
| RelA | Cell Signaling | CST 8242 | D14E12 | Rabbit | 1:1000 |

### 7. Apoptosis and Matrigel Tube Formation Assays

Assays were conducted according to the manufacturer’s instructions and as previously described [60, 61]. Tube formation was assessed with the Angiogenesis Analyzer package in ImageJ [62].

### 8. RNA Sequencing

Human PAECs were grown to 80% confluency on gel coated plates and transfected with either *Fyn*, *Lck*, or nontargeted siRNA. At 48 hours, cells were placed in 10% EGM starvation media. Two hours prior to harvest all cells were treated with either PBS in 10% EGM or, in one case, 20 ng/mL of BMP9 in 10% EGM. Cells were harvested and mRNA was extracted via Qiagen mini columns. RNA quality was assessed with the Agilent 2100 Bioanalyzer system and found to have a RIN of 9.8-9.9 (Supplemental Figure 5-A). cDNA libraries were generated (Illumina), barcoded, and sequenced via a paired-end 150 bp sequencing strategy to a depth of between 38 and 46 million paired-end reads on the Illumina platform. Raw reads were mapped to the human genome using STAR [63]. Quality control metrics regarding sequencing and alignment are given in Supplemental Figure 5-A. Differential gene expression analysis was performed using the R package DEseq2 [64]. Briefly, differentially expressed genes were calculated via the “DEseq” command after setting transfection condition (FYN, LCK, NT) and treatment (PBS, BMP9) as factors. Conditions were contrasted against the nontargeted PBS condition to determine the fold change and adjusted p-value for each gene in each condition. A variance stabilizing transformation of the raw count data was used to generate the gene expression heat map. Next, differentially expressed genes present in each condition were found using a cutoff Log2 fold change of 0.3 (fold change of less than 0.8 or greater than 1.25) with a false discovery rate of 5% and the R function “venn.diagram”. Lists of significant differentially expressed genes that were present in select conditions (IE increased in the LCK case and decreased in the FYN case) were used for gene ontology analysis. Gene ontology and transcription factor analysis was facilitated using the Enrichr GSEA web server [34, 65]. For transcription factor analysis, the packages TRRUST, BART, and oPOSSUM-3 [35–37] were used to predict transcription factors co-regulating the genes in query set. Results were tailed to show which transcription factors were predicted by different methods.

### 9. Analysis of Public PBMC Single Cell RNA Seq Data

Raw count data of 2700 PBMCs (PBMCs from a health human donor, single cell immune profiling dataset by Cell Ranger 1.1.0 on Illumina NextSeq 500 with ∼69,000 reads per cell, 10x Genomics, 2016, May 26, published in [29]) were analyzed using the Seurat V 3.0 sc-RNA seq pipeline to perform unsupervised hierarchical clustering [66]. Average gene expression was calculated on a per-cluster basis. Cell identity was determined based on differentially upregulated genes in a cluster of interest relative to all the other clusters in the dataset (Supplemental Figure 2). After clusters were developed and identified, the average expression of genes of interest (*Bmpr2, Id1, Id2, Id3, Lck)* was determined. The percent of cells expressing the gene of interest (defined as a cell with > 1 count) was also calculated. Both the magnitude of expression (presented as a Z-score relative to all cells in the sample) and the percent expression are presented in dot plot format.

### 10. Immunofluorescence Staining of Adherent Cells and Human Lung Tissues

For confocal localization of proteins in cells, cells were washed in warm PBS and then fixed in 10% normal buffered formalin for 20 minutes at room temperature. Cells were washed with PBS, then permeabilized and blocked with 0.3% Triton X-100 and 5% Serum for 1-3 hours at room temperature. Primary antibodies were incubated overnight at 4C at the following dilutions in a 5% serum buffer in PBS: LCK (ABCAM Ab227975, 1:200), FYN (BD Biosciences BD61063 1:150), phopho-P65 RelA (Cell Signaling Technologies 93H1,1:200). Appropriate secondary antibodies were used at a 1:250 dilution in a 5% serum PBS buffer for 1 hour. Cells were washed in PBS + 0.5% Tween-20 x 3. In all cases DAPI counter stains or F-Actin counter stains (fluorescently labeled phalloidin) were used per published protocols. Cells were imaged by confocal microscopy. Quantification of signal was performed in ImageJ.

### 11. Single Molecule mRNA Fluorescent in situ Hybridization (sm-FISH)

Sm-FISH for *Lck* was performed using the RNA-scope protocol [67] (Advanced Cell Diagnostics, Inc.) using the Channel 1 probe to *Lck* (Cat No. 440201). After completion of the sm-FISH protocol, tissue was counter stained for endothelial (VE-Cadherin, R&D AF938, 1:300) and smooth muscle (Acta2, Direct a488 Conjugation, 1A4 Clone, 1:200). Tissue was imaged by confocal microscopy and analyzed with ImageJ.

### 12. Statistical Analysis

Data were analyzed using R/R-Studio version 1.4.1106. Statistical tests were performed as appropriate and included the following: Student’s t-test, One-Way ANOVA and Two-way ANOVA, followed by the appropriate post-hoc test, as indicated. Bars show mean ± Standard Deviation. Differences are considered statistically significant for p values less than 0.05. All significant p values are reported on graphs.

### 13. Drug Prediction by Broad Institute Connectivity Map

The Connectivity Map (CMap) portal (https://clue.io) was used to perform all searches. First, the genes returned by RNA seq were narrowed to only include CMap “Best INFerred” or BING genes. These are genes used in CMap that are computationally inferred with high confidence from the 978 genes actively profiled by the L1000 platform [41]. Next, the differentially expressed genes were ranked based on the overall magnitude of change, such that:

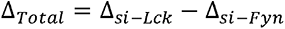

Genes were ranked based on Δ*_Total_*. Gene with a positive Δ*_Total_* (e.g. increased with si-Lck, decreased with si-Fyn) comprised the “Inflammatory” signature whereas genes with a negative Δ*Total* comprised the “endothelial” signature. Genes with the highest |Δ_*Total*_| were filtered against the list of available CMap BING genes. The remaining BING gene-based angiogenic and inflammatory signatures were truncated to lists of lengths 10, 15, 20, 25, 50, 75, 100, 125 and 150 genes long. Each gene list (both angiogenic and inflammatory, n = 18) was then submitted in the CMap “Query” program as an Up-Regulated gene. Query parameters are “Gene expression (L1000)”, “Touchstone”, “Individual Query”, and “Latest.” All searches were performed between 11/2021 and 12/2021. The search results were opened online in CMap Morpheus, and then downloaded as a GCT version 1.3 file from the “query_result” tab. A custom R script was used to open the 18 GCT files and combine them into a single data structure that can be subset by gene signature, gene search length, perturbagen class, search query correlation score, cell type, transcriptional activation score (TAS), and false discovery rate. This data set was used to generate Figure 6.

Regarding number of genes used in CMap search query. Aggregate search results are shown in Sup. Figure 6A. There was a linear relationship between the number of genes used in the search and the number of significant perturbagens (defined as having a false discovery rate of < 5%) returned by querying the inflammatory signature. The endothelial signature exhibited a diminishing return of perturbagens after 75 genes were used. Going forward, we used the 150-gene inflammatory signature search and the 75-gene endothelial signature search because they returned the most perturbagens.

To determine which cell lines are acceptable substrates for PAEC drug prediction, we leveraged the fact that multiple CMap cell lines were transcriptionally profiled after undergoing either knockout or overexpression of *FYN* and *LCK*. CMap gene manipulations occur via short hairpin (SH) RNA knockdown, plasmid overexpression (OE), and Crisper Cas9 induced loss of function. We reasoned that cell lines in which the CMap sh-*LCK* or sh-*FYN* perturbagen positively connects with our own PAEC *LCK* or *FYN* knockout gene signature, or where the CMap OE-*LCK* and OE-*FYN* perturbagen negatively connects with our PAEC *LCK* or *FYN* knockout gene signature would be the best cell substrates for drug prediction. Results of this analysis are shown in Sup. Figure 6B-C.

To determine which drugs would be used for preliminary testing, we used the following criteria: Candidate drugs should be (1) a TKI, (2) located in the periphery of Figure 6E (highest magnitude of endothelial and inflammatory connectivity scores), (3) have low FDR and high transcriptional activation scores (TAS) scores, and (4) are commercial availability. Note that TAS is a measure of signature strength and replicate correlation intended to represent a perturbagen’s transcriptional activity. The most promising candidates from preliminary screening were selected from each quadrant. The ten candidate drugs included Brivanib, JW-55, Ulixertinib, Pazopanib, Regorafenib, Alvocidib, TAK-715, H-9 Dihydrochloride, Mycophenolic acid, and BMS 345541.

### 14. Drug Treatment in Cell Lines

Human PAECs were grown to near confluency (90%) and used between passages 4-6. All drugs were diluted in DMSO at concentrations like those used in the CMap/L1000 experiments in HUVEC cell lines. Drug dilutions were prepared in DMSO and added to full endothelial growth media (EGM). Final concentrations were: Brivanib (10 μM), Quizartinib (10 μM), Regorafanib (37 μM), and Pazopanib (37 μM). Cells were starved for 6 hours in 10% EGM prior to giving drug or DMSO. Drug/DMSO in full EGM was added to cells for a total treatment time of 24 hours. For Brivanib and Quizartinib, either BMP9 at 10 ng/mL or PBS was added 2 hours prior to harvesting the cells. Cells were then harvested, run on SDS-Page, and immunoblotted for Id1, pSMAD1, and pSMAD3. For Regorafenib and Pazopanib, cells were grown in gelatin coated slide well chambers then treated with drug for 24 hours. Each slide contained a drug replicate and a DMSO only replicate. After 24 hours, slides were fixed in 10% NBF for 25 minutes, then blocked with 5% goat serum, 1% BSA, and 0.3% Triton in PBS for 1 hour prior to incubating with p65-RelA primary antibody and Alexa-488 conjugated phalloidin overnight in the same buffer. Cells were treated with DAPI and an Alexa-633 conjugated goat anti-rabbit secondary antibody at a dilution of 1:250 for 1 hour. Slides were then imaged by confocal microscopy, collecting z-stacks at Nyquist spacing through the entire cell biolayer. Between 4 and 6 images were collected of 4 experiments each. Images were analyzed with Imaris 3D processing software. DAPI staining was used to define a 3D volume representing the nucleus. To control for variations in staining, the DAPI intensity threshold at which to call a given voxel intranuclear was adjusted on a per-experiment basis. This ensured that the average nuclear diameter for all baseline DMSO experiments were the same. P65 RelA signal intensity was quantified on a per voxel basis for each voxel contained in the DAPI nucleus. This data was used to compute the average p65-RelA expression for the nuclear volume for each intact nucleus imaged. Average p65-RelA intensity and overall nuclear size for Pazopanib and Regorafenib treated cells were compared relative to their DMSO counterparts which were stained on the same slide with the same primary and secondary antibody buffers.

## Author Contributions

AMA and ES conceived of experiments and study design, revised the manuscript. AMA did all experiments, RNA sequencing and downstream analysis, performed data analysis, performed statistical analysis, and wrote the manuscript. XT conducted the high throughput screen. MKA assisted with cell culture.

## Acknowledgements

We thank Maya Kumar for her assistance in reviewing the text.

## Abbreviations

BMPR2: Bone Morphogenetic Protein Receptor Type 2
BRE: BMP Response Element
CMap: Broad Institute Connectivity Map Database
DEG: Differentially Expressed Gene
EC: Endothelial Cell
ECD: Endothelial Cell Dysfunction
FYN: FYN Proto-Oncogene
HTS: High Throughput siRNA Screen
ID1,2, and 3: Inhibitor of Differentiation 1, 2, and 3
LCK: Lymphocyte specific protein tyrosine kinase
LINCS/L1000: Library of Integrated Network-Based Cellular Signatures
OE: Overexpression
PAEC: Pulmonary Artery Endothelial Cell
PAH: Pulmonary Arterial Hypertension
PBMC: Peripheral Blood Mononuclear Cell
PDGFR: Platelet derived growth factor receptor
pSMAD1: Phosphorylated SMAD-1
PBS: Phosphate Buffered Saline
PTK: Protein Tyrosine Kinase
ROS: Reactive Oxygen Species
scRNA-seq: Single Cell RNA Seq
SH: Short Hairpin RNA
siRNA: Small Interfering RNA
si-Fyn: PAECs subject to *Fyn* knockout with si-RNA
si-Lck: PAECs subject to *Lck* knockout with si-RNA
SRC: *Src-*Proto-Oncogene, Non-Receptor Tyrosine Kinase
SrcA: SRC-Family A Protein Tyrosine Kinase
SrcB: SRC-Family B Protein Tyrosine Kinase
TCR: T-Cell Receptor
TKI: Tyrosine Kinase Inhibitor
VEGFR: Vascular Endothelial Growth Factor Receptor

---

**Supplemental Table 1:** Baseline characteristics and clinical data of endothelial cells extracted from the lungs of 3 PAH patients and 3 failed lung transplant donor lungs grown in tissue culture and assessed by western blot in Figure 2.

| Lane | ID | Donor/Patient | Etiology | Age | Sex | Race | 6 Minute Walk<br>Test Distance | Therapy |
| --- | --- | --- | --- | --- | --- | --- | --- | --- |
|  |  |  |  | Years |  |  | Ft. (m) |  |
| 1 | DC-1 | Donor Control | Stoke, ICH | 34 | Female | Asian | - | - |
| 2 | DC-2 | Donor Control | Acute MI | 57 | Female | White | - | - |
| 3 | DC-3 | Donor Control | Blunt Trauma | 33 | Female | White | - | - |
| 4 | FPAH-1 | Patient | HPAH | 22 | Female | White | 1233 (375.8) | PDE5i, ERA, PC |
| 5 | FPAH-2 | Patient | HPAH | 27 | Female | White | 1180 (359.7) | PDE5i, ERA, PC |
| 6 | IPAH | Patient | IPAH | 39 | Female | Latinx | 860 (262.1) | PDE5i, ERA, PC |

PAH = Pulmonary Arterial Hypertension
HPAH = Hereditary Pulmonary Arterial Hypertension
IPAH = Idiopathic Pulmonary Arterial Hypertension
PDE5i = Phosphodiesterase type 5 Inhibitor
ERA = Endothelin receptor antagonist
PC = Prostacyclin

**Supplemental Figure 1:**
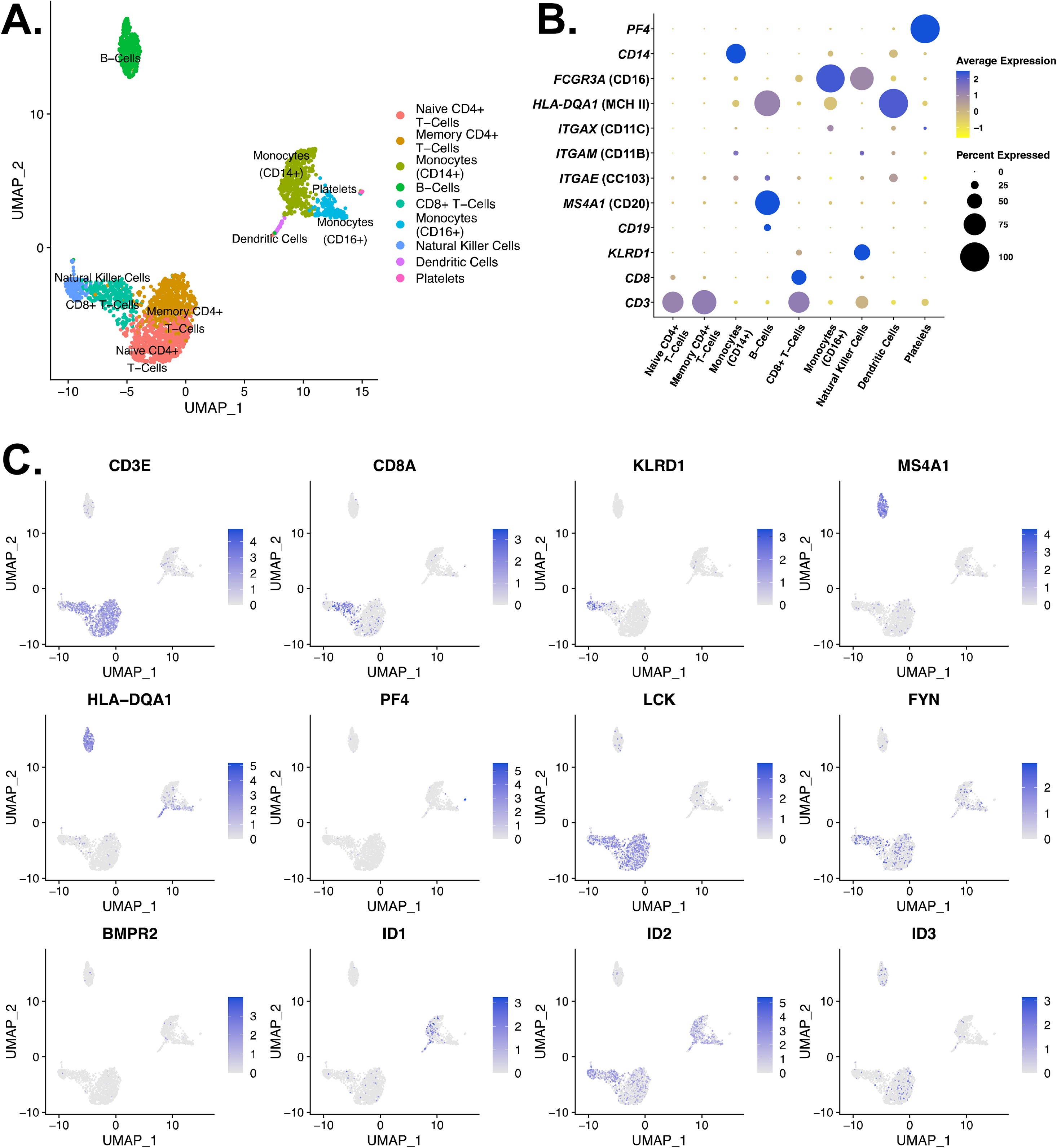
**(A)** UMAP plot of all 2700 human PBCMs after unsupervised clustering by the Seurat single cell RNA seq pipeline. Nine distinct clusters are identified. Clusters have been annotated for cell identity based on the gene expression detailed in **(2 B-C)**. **(B)** Dot plots showing expression of canonical genes known to be expressed in specific cell types which informs cluster identity. *CD3* denotes T-cells, *CD8* indicates CD8+ T-Cells, *KLRD1* (CD94) denotes natural killer cells, *CD19* and *MS4A1* (CD20) denotes B- Cells, *ITGAE* (Cd103)*, HLA-CQA1* (MHC II), and *ITGAX* (Cd11c) denote dendritic cells, FCGR3A (Cd16) and *CD14* denotes monocytes, and *PF4* indicates platelets. Dot size indicates percent expression of the gene (the number of cells in the cluster that have > 1 counts of a given gene). Average expression is the Z-score of the log-normalized counts of a gene within the cluster. **(C)** Feature plots of all genes of interested examined in PBMCs for correlation with the clusters of Supplemental Figure 1-A.

**Supplemental Figure 2:**
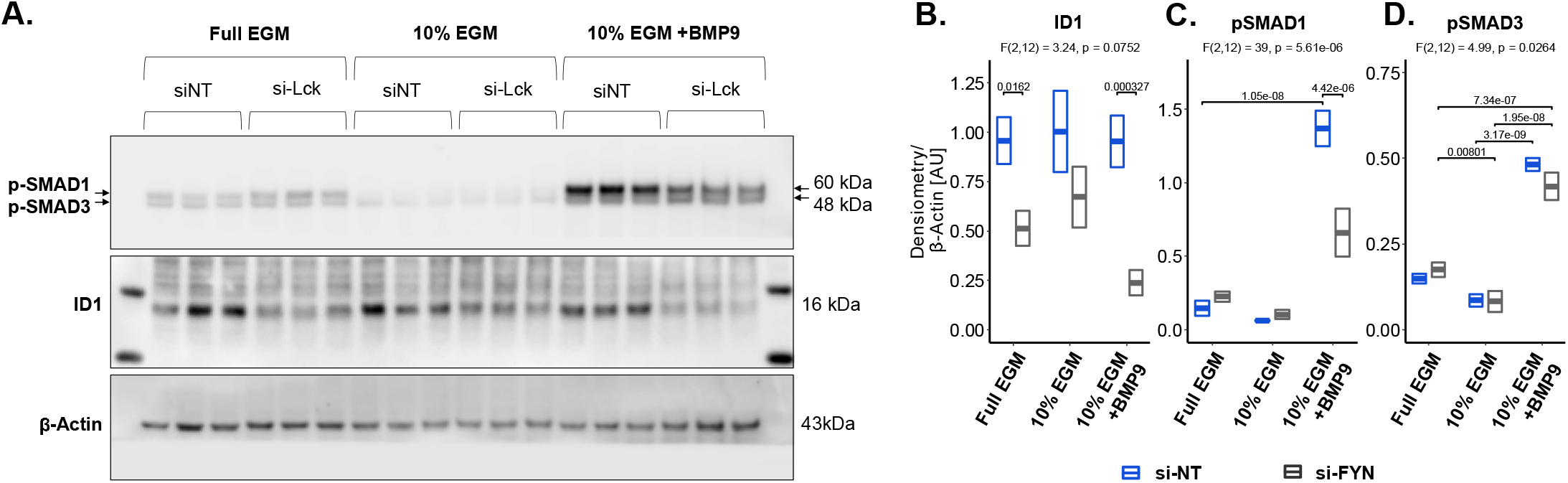
**(A)** Knockout of *LCK* in PAECs suppresses canonical BMPR2 signaling in different media conditions. PAECs were transfected with siRNA targeting *LCK* (si-LCK) or a nontargeting siRNA (siNT) for 6 hours. 24 hours after transfection, the cells were either starved in 10% endothelial growth media (EGM) or kept in full EGM. 1.5 hours prior to cell lysis, the starvation cells were either treated with PBS or 20 ng/mL of BMP9. Cell lysate was run on SDS-PAGE. **(B-D)** Phospho-SMAD1 was significantly upregulated by BMP9 treatment and was significantly reduced in the si-LCK group. Id1 was significantly reduced by siLck in both the full EGM and the BMP9 case. 2-Way ANOVA for the interaction between treatment (Full EGM, 10% EGM, 10% EGM + BMP9) and transfection (si-LCK and si-NT) with Tukey’s Range Test was used to determine significance. Normality checks for ANOVA residuals and Levene’s test for homogeneity were carried out and assumptions met.

**Supplemental Figure 3:**
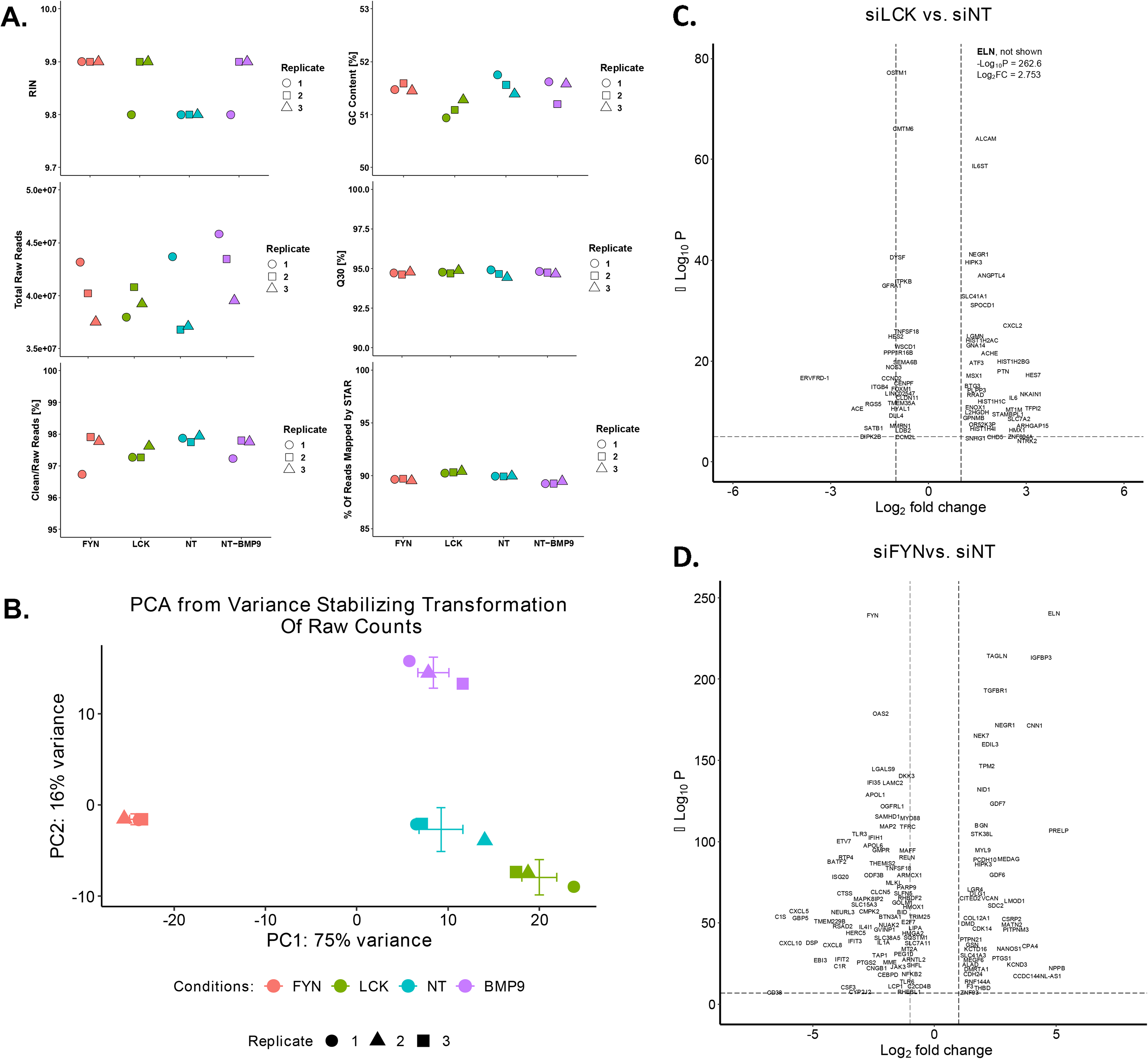
Quality control metrics and volcano plots for the RNA seq analysis detailed in Figure 5. **(A)** Shows the distribution of RIN, Total Raw Reads, Clean/Raw Reads [%], GC Content [%], Q30 [%], and % of Reads Mapped by STAR. **(B)** Principal component analysis of all samples used in analysis shows that all conditions separate with replicates clustering together. Detailed volcano plot of differentially expressed genes in the **(C)** siLCK and **(D)** siFYN differential gene expression relative to the siNT case. Note that *ELN* is not depicted to allow a better view of all genes on the si-LCK plot.

**Supplemental Figure 4:**
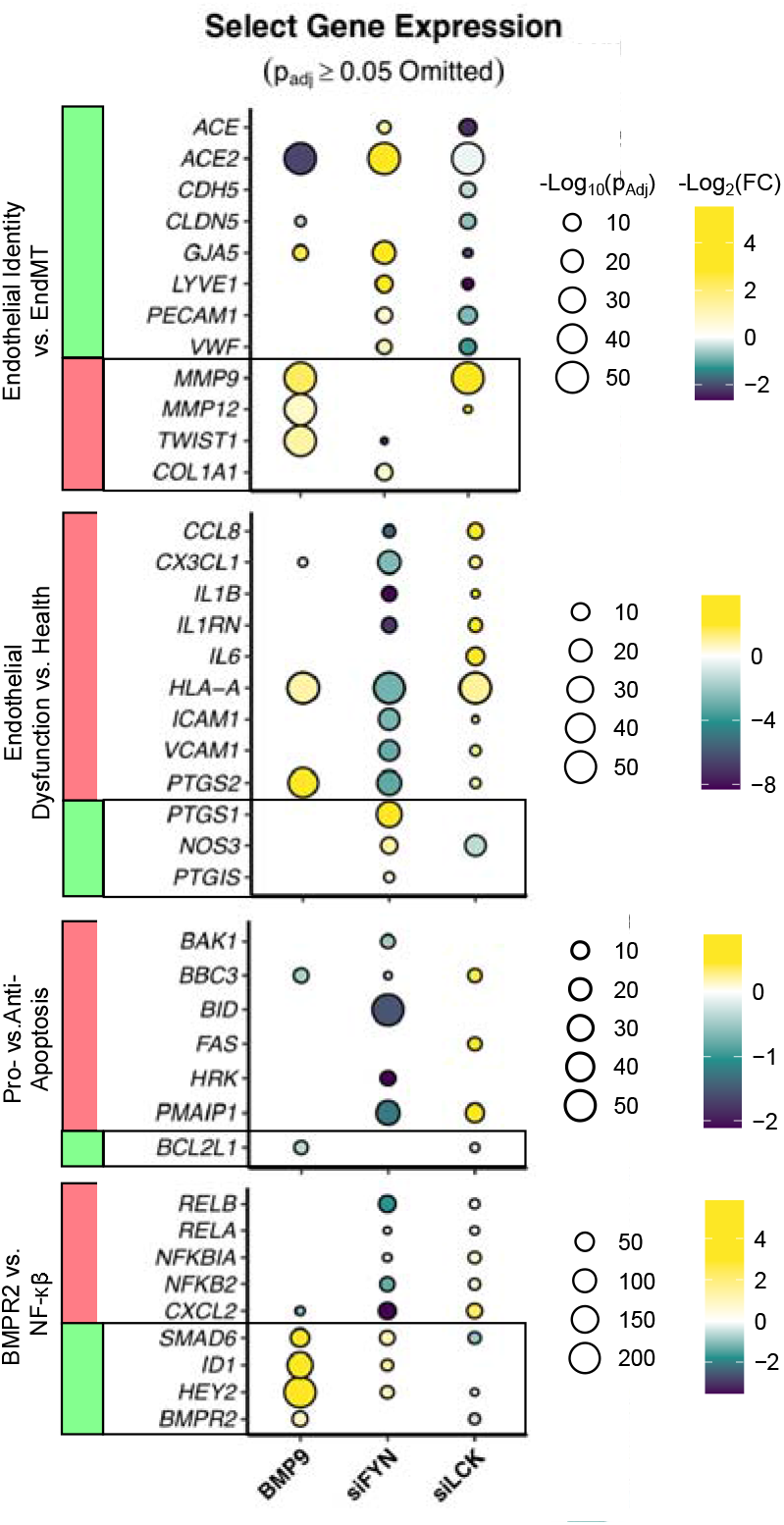
A biased assessment of endothelial specific genes in the RNA seq analysis further supports a differential effect of *FYN* and *LCK* knockout in human pulmonary artery endothelial cells with respect to apoptosis and endothelial cell dysfunction. We find a loss of canonical endothelial markers (*GJA5, LYVE1, VWF, PECAM1*). Additionally, genes for endothelial specific functions such as nitric oxide synthase (*NOS3*) are decreased by *LCK* knockout and increased by *FYN* knockout. *FYN* knockout is also associated with increased prostaglandin synthase 1 (*PTGS1*) expression.

**Supplemental Figure 5:**
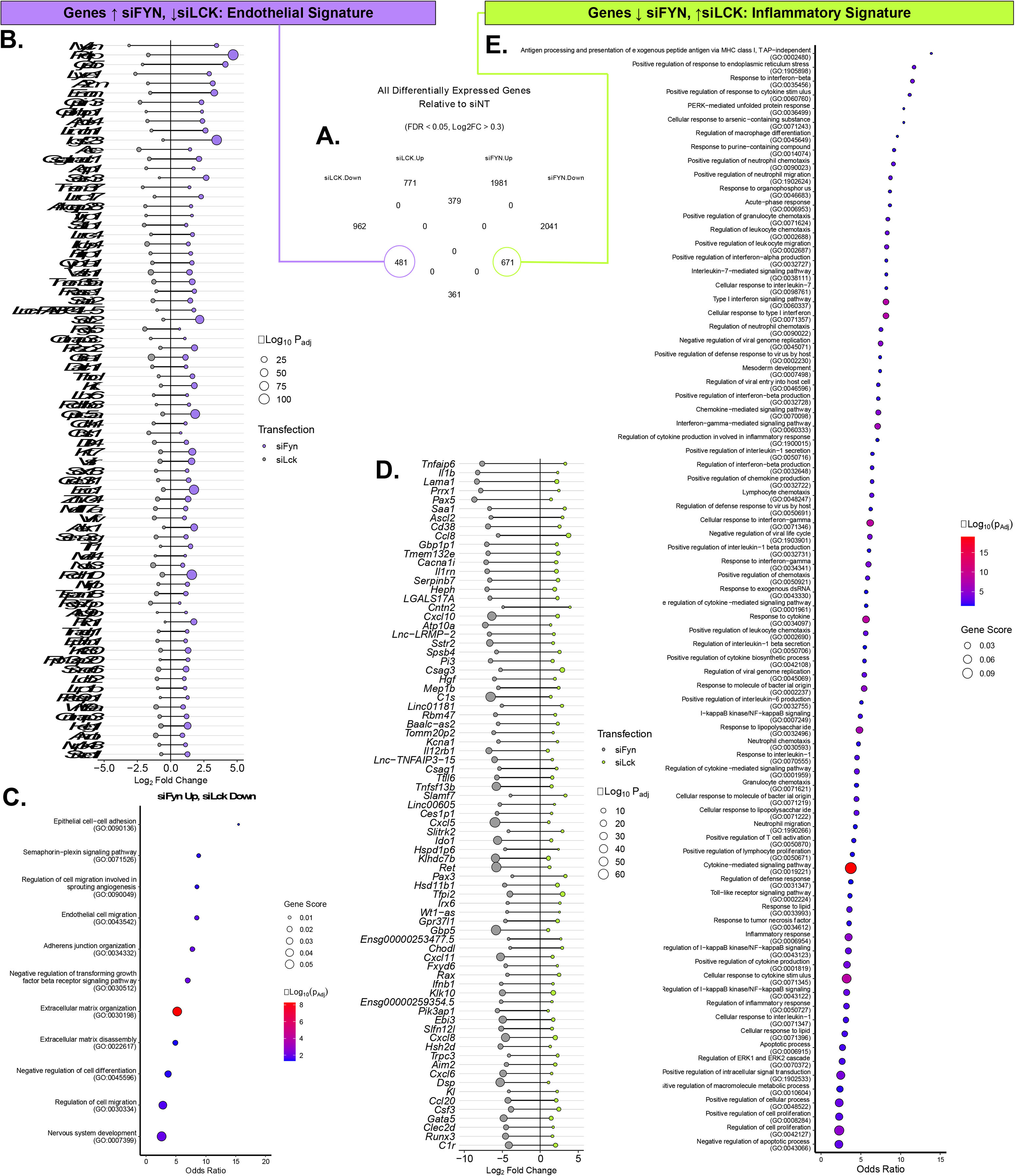
“Inflammatory” and “Endothelial” gene signatures. Extended list of genes co- differentially expressed and are increased by si-LCK and decreased si-FYN. Because si-LCK imparts a dysfunctional EC phenotype, genes in this list are implicated in EC dysfunction. Genes in this list are increased by *LCK* suppression and decreased by *FYN* suppression. The accompanying gene ontology analysis shows that multiple pathways are implicated that represent a response to interferon signaling, lipopolysaccharide, NF-κB signaling, or neutrophil transmigration.

**Supplemental Figure 6:**
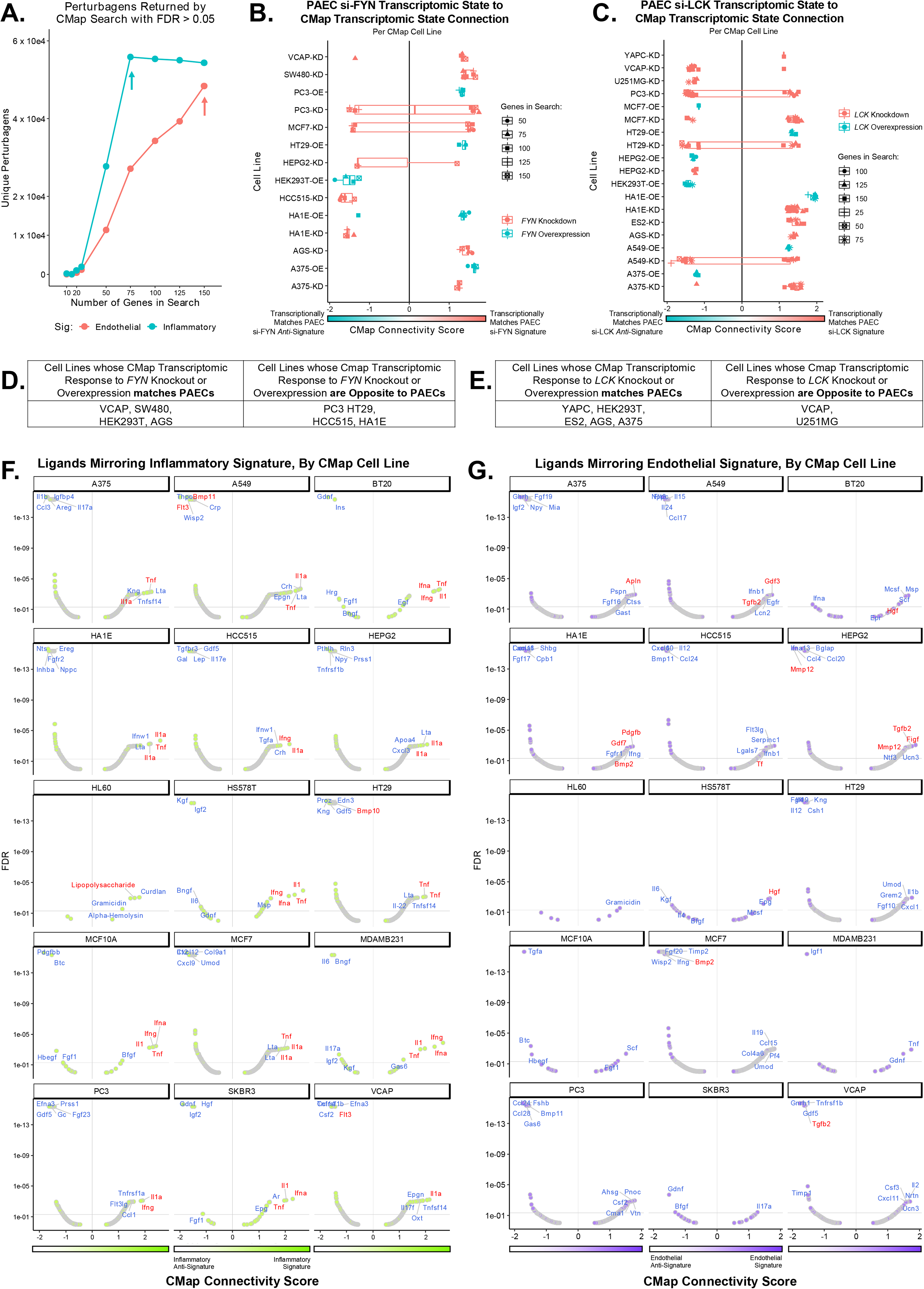
Extension of Figure 6C: Full results for Connectivity Map (CMap) analysis detailing ligands which recapitulate the inflammatory and endothelial signatures. Data for all cell lines present in CMap data is shown. Y axis indicates significance by -Log10(False Discovery Rate) and X axis indicates the Connectivity Score.

**Supplemental Figure 7:**
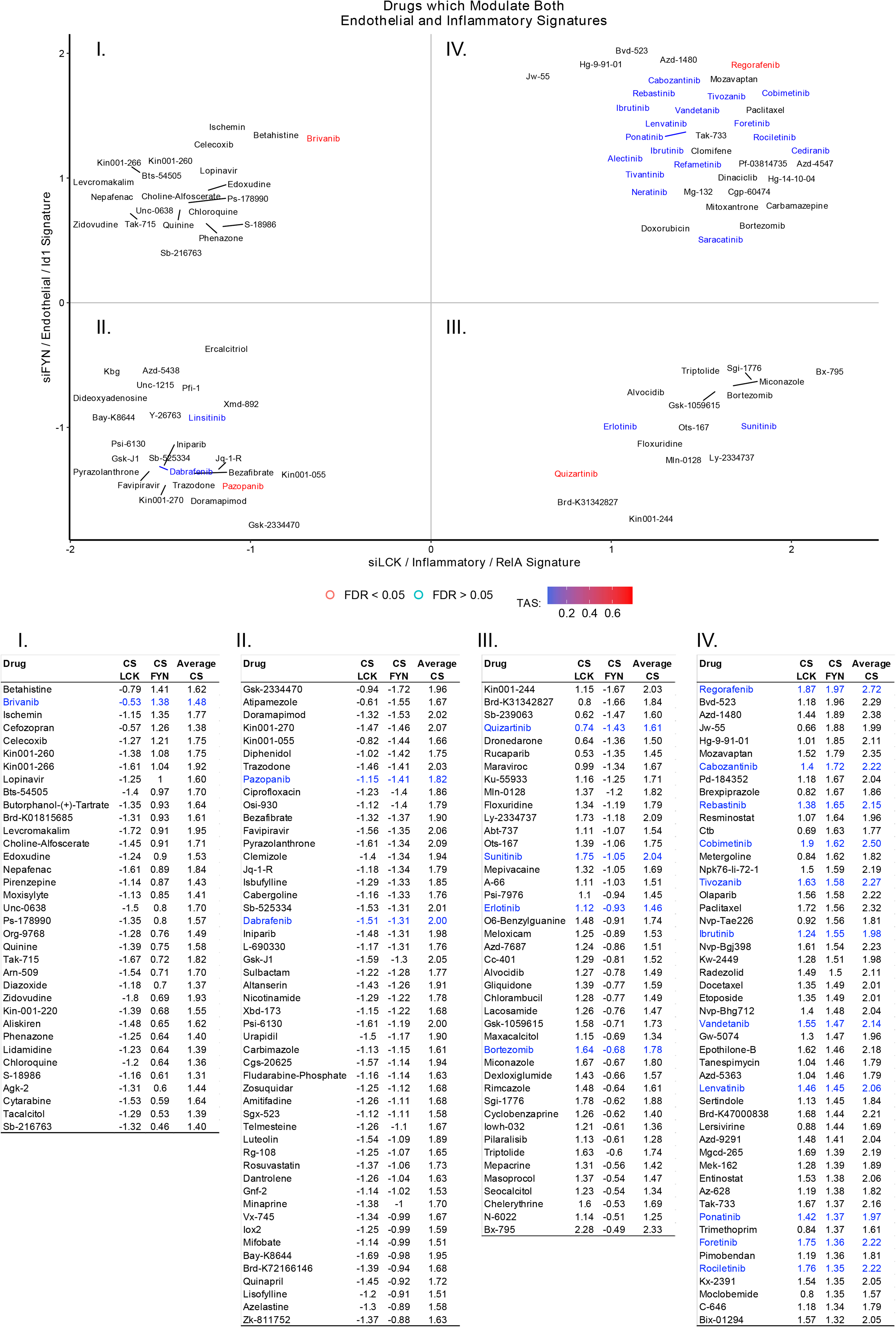
Extended data of CMap search results.

## References Cited

1. G. Manning DBW, R. Martinez, T. Hunter and S. Sudarsanam. The Protein Kinase Complement of the Human Genome. Science. 2002;298(5600):1912-6+33-34.

2. Grimminger F, Schermuly RT, Ghofrani HA. Targeting non-malignant disorders with tyrosine kinase inhibitors. Nat Rev Drug Discov. 2010;9(12):956–70. Epub 2010/12/02. doi: 10.1038/nrd3297. PubMed PMID: 21119733.

3. Rubin LJ. Primary pulmonary hypertension. N Engl J Med. 1997;336(2):111–7. Epub 1997/01/09. doi: 10.1056/NEJM199701093360207. PubMed PMID: 8988890.

4. Machado RD, Eickelberg O, Elliott CG, Geraci MW, Hanaoka M, Loyd JE, Newman JH, Phillips JA, 3rd, Soubrier F, Trembath RC, Chung WK. Genetics and genomics of pulmonary arterial hypertension. J Am Coll Cardiol. 2009;54(1 Suppl):S32-42. doi: 10.1016/j.jacc.2009.04.015. PubMed PMID: 19555857; PMCID: PMC3725550.

5. Rabinovitch M. Molecular pathogenesis of pulmonary arterial hypertension. J Clin Invest. 2012;122(12):4306–13. doi: 10.1172/JCI60658. PubMed PMID: 23202738; PMCID: PMC3533531.

6. Lau EMT, Giannoulatou E, Celermajer DS, Humbert M. Epidemiology and treatment of pulmonary arterial hypertension. Nat Rev Cardiol. 2017;14(10):603–14. doi: 10.1038/nrcardio.2017.84. PubMed PMID: 28593996.

7. Hoeper MM, Barst RJ, Bourge RC, Feldman J, Frost AE, Galie N, Gomez-Sanchez MA, Grimminger F, Grunig E, Hassoun PM, Morrell NW, Peacock AJ, Satoh T, Simonneau G, Tapson VF, Torres F, Lawrence D, Quinn DA, Ghofrani HA. Imatinib mesylate as add-on therapy for pulmonary arterial hypertension: results of the randomized IMPRES study. Circulation. 2013;127(10):1128–38. doi: 10.1161/CIRCULATIONAHA.112.000765. PubMed PMID: 23403476.

8. Pankey EA, Thammasiboon S, Lasker GF, Baber S, Lasky JA, Kadowitz PJ. Imatinib attenuates monocrotaline pulmonary hypertension and has potent vasodilator activity in pulmonary and systemic vascular beds in the rat. Am J Physiol Heart Circ Physiol. 2013;305(9):H1288-96. doi: 10.1152/ajpheart.00329.2013. PubMed PMID: 23997103; PMCID: PMC3840242.

9. Rasheed W, Flaim B, Seymour JF. Reversible severe pulmonary hypertension secondary to dasatinib in a patient with chronic myeloid leukemia. Leuk Res. 2009;33(6):861–4. doi: 10.1016/j.leukres.2008.09.026. PubMed PMID: 18986702.

10. Montani D, Bergot E, Gunther S, Savale L, Bergeron A, Bourdin A, Bouvaist H, Canuet M, Pison C, Macro M, Poubeau P, Girerd B, Natali D, Guignabert C, Perros F, O’Callaghan DS, Jais X, Tubert-Bitter P, Zalcman G, Sitbon O, Simonneau G, Humbert M. Pulmonary arterial hypertension in patients treated by dasatinib. Circulation. 2012;125(17):2128–37. doi: 10.1161/CIRCULATIONAHA.111.079921. PubMed PMID: 22451584.

11. Guignabert C, Phan C, Seferian A, Huertas A, Tu L, Thuillet R, Sattler C, Le Hiress M, Tamura Y, Jutant EM, Chaumais MC, Bouchet S, Maneglier B, Molimard M, Rousselot P, Sitbon O, Simonneau G, Montani D, Humbert M. Dasatinib induces lung vascular toxicity and predisposes to pulmonary hypertension. J Clin Invest. 2016;126(9):3207–18. doi: 10.1172/JCI86249. PubMed PMID: 27482885; PMCID: PMC5004960.

12. Hickey PM, Thompson AA, Charalampopoulos A, Elliot CA, Hamilton N, Kiely DG, Lawrie A, Sabroe I, Condliffe R. Bosutinib therapy resulting in severe deterioration of pre-existing pulmonary arterial hypertension. Eur Respir J. 2016;48(5):1514–6. doi: 10.1183/13993003.01004-2016. PubMed PMID: 27660511.

13. Quilot FM, Georges M, Favrolt N, Beltramo G, Foignot C, Grandvuillemin A, Montani D, Bonniaud P, Camus P. Pulmonary hypertension associated with ponatinib therapy. Eur Respir J. 2016;47(2):676–9. doi: 10.1183/13993003.01110-2015. PubMed PMID: 26743481.

14. Chabrol A, Mayenga M, Hamid AM, Friard S, Salvator H, Doubre H, Fraboulet S, Metivier AC, Catherinot E, Rivaud E, Chaumais MC, Montani D, Couderc LJ, Tcherakian C. Lorlatinib - Induced pulmonary arterial hypertension. Lung Cancer. 2018;120:60–1. Epub 2018/05/12. doi: 10.1016/j.lungcan.2018.03.023. PubMed PMID: 29748016.

15. Swords R, Mahalingam D, Padmanabhan S, Carew J, Giles F. Nilotinib: optimal therapy for patients with chronic myeloid leukemia and resistance or intolerance to imatinib. Drug Des Devel Ther. 2009;3:89–101. Epub 2009/11/19. doi: 10.2147/dddt.s3069. PubMed PMID: 19920925; PMCID: PMC2769239.

16. Gover-Proaktor A, Granot G, Pasmanik-Chor M, Pasvolsky O, Shapira S, Raz O, Raanani P, Leader A. Bosutinib, dasatinib, imatinib, nilotinib, and ponatinib differentially affect the vascular molecular pathways and functionality of human endothelial cells. Leuk Lymphoma. 2019;60(1):189–99. Epub 2018/05/10. doi: 10.1080/10428194.2018.1466294. PubMed PMID: 29741440.

17. Dasgupta SK, Le A, Vijayan KV, Thiagarajan P. Dasatinib inhibits actin fiber reorganization and promotes endothelial cell permeability through RhoA-ROCK pathway. Cancer Med. 2017;6(4):809–18. Epub 2017/03/21. doi: 10.1002/cam4.1019. PubMed PMID: 28316141; PMCID: PMC5387130.

18. Phan C, Jutant EM, Tu L, Thuillet R, Seferian A, Montani D, Huertas A, Bezu JV, Breijer F, Vonk Noordegraaf A, Humbert M, Aman J, Guignabert C. Dasatinib increases endothelial permeability leading to pleural effusion. Eur Respir J. 2018;51(1). Epub 2018/01/20. doi: 10.1183/13993003.01096-2017. PubMed PMID: 29348177.

19. Cornet L, Khouri C, Roustit M, Guignabert C, Chaumais MC, Humbert M, Revol B, Despas F, Montani D, Cracowski JL. Pulmonary arterial hypertension associated with protein kinase inhibitors: a pharmacovigilance-pharmacodynamic study. Eur Respir J. 2019;53(5). doi: 10.1183/13993003.02472-2018. PubMed PMID: 30846469.

20. Zhu N, Gonzaga-Jauregui C, Welch CL, Ma L, Qi H, King AK, Krishnan U, Rosenzweig EB, Ivy DD, Austin ED, Hamid R, Nichols WC, Pauciulo MW, Lutz KA, Sawle A, Reid JG, Overton JD, Baras A, Dewey F, Shen Y, Chung WK. Exome Sequencing in Children With Pulmonary Arterial Hypertension Demonstrates Differences Compared With Adults. Circ Genom Precis Med. 2018;11(4):e001887. doi: 10.1161/CIRCGEN.117.001887. PubMed PMID: 29631995; PMCID: PMC5896781.

21. Andruska A, Spiekerkoetter E. Consequences of BMPR2 Deficiency in the Pulmonary Vasculature and Beyond: Contributions to Pulmonary Arterial Hypertension. Int J Mol Sci. 2018;19(9). doi: 10.3390/ijms19092499. PubMed PMID: 30149506.

22. Frump AL, Lowery JW, Hamid R, Austin ED, de Caestecker M. Abnormal trafficking of endogenously expressed BMPR2 mutant allelic products in patients with heritable pulmonary arterial hypertension. PLoS One. 2013;8(11):e80319. doi: 10.1371/journal.pone.0080319. PubMed PMID: 24224048; PMCID: PMC3818254.

23. Prewitt AR, Ghose S, Frump AL, Datta A, Austin ED, Kenworthy AK, de Caestecker MP. Heterozygous null bone morphogenetic protein receptor type 2 mutations promote SRC kinase-dependent caveolar trafficking defects and endothelial dysfunction in pulmonary arterial hypertension. J Biol Chem. 2015;290(2):960–71. doi: 10.1074/jbc.M114.591057. PubMed PMID: 25411245; PMCID: PMC4294523.

24. Dannewitz Prosseda S, Tian X, Kuramoto K, Boehm M, Sudheendra D, Miyagawa K, Zhang F, Solow-Cordero D, Saldivar JC, Austin ED, Loyd JE, Wheeler L, Andruska A, Donato M, Wang L, Huebner K, Metzger RJ, Khatri P, Spiekerkoetter E. Fragile Histidine Triad (FHIT), a Novel Modifier Gene in Pulmonary Arterial Hypertension. Am J Respir Crit Care Med. 2018. doi: 10.1164/rccm.201712-2553OC. PubMed PMID: 30107138.

25. 25. Zilberberg L, ten Dijke P, Sakai LY, Rifkin DB. A rapid and sensitive bioassay to measure bone morphogenetic protein activity. BMC Cell Biol. 2007;8:41. Epub 2007/09/21. doi: 10.1186/1471-2121-8-41. PubMed PMID: 17880711; PMCID: PMC2094707.

26. Pullamsetti SS, Berghausen EM, Dabral S, Tretyn A, Butrous E, Savai R, Butrous G, Dahal BK, Brandes RP, Ghofrani HA, Weissmann N, Grimminger F, Seeger W, Rosenkranz S, Schermuly RT. Role of Src tyrosine kinases in experimental pulmonary hypertension. Arterioscler Thromb Vasc Biol. 2012;32(6):1354–65. doi: 10.1161/ATVBAHA.112.248500. PubMed PMID: 22516066.

27. Kim DJ, Norden PR, Salvador J, Barry DM, Bowers SLK, Cleaver O, Davis GE. Src- and Fyn-dependent apical membrane trafficking events control endothelial lumen formation during vascular tube morphogenesis. PLoS One. 2017;12(9):e0184461. doi: 10.1371/journal.pone.0184461. PubMed PMID: 28910325; PMCID: PMC5598984.

28. Baumgart B, Guha M, Hennan J, Li J, Woicke J, Simic D, Graziano M, Wallis N, Sanderson T, Bunch RT. In vitro and in vivo evaluation of dasatinib and imatinib on physiological parameters of pulmonary arterial hypertension. Cancer Chemother Pharmacol. 2017;79(4):711–23. doi: 10.1007/s00280-017-3264-2. PubMed PMID: 28283735.

29. Butler A, Hoffman P, Smibert P, Papalexi E, Satija R. Integrating single-cell transcriptomic data across different conditions, technologies, and species. Nature Biotechnology. 2018;36(5):411–20. doi: 10.1038/nbt.4096.

30. Richard KC, Bertolesi GE, Dunfield LD, McMaster CR, Nachtigal MW. TSAd interacts with Smad2 and Smad3. Biochem Biophys Res Commun. 2006;347(1):266–72. doi: 10.1016/j.bbrc.2006.06.068. PubMed PMID: 16806069.

31. David L, Mallet C, Mazerbourg S, Feige JJ, Bailly S. Identification of BMP9 and BMP10 as functional activators of the orphan activin receptor-like kinase 1 (ALK1) in endothelial cells. Blood. 2007;109(5):1953–61. doi: 10.1182/blood-2006-07-034124. PubMed PMID: 17068149.

32. Gimbrone MA, Jr., Garcia-Cardena G. Endothelial Cell Dysfunction and the Pathobiology of Atherosclerosis. Circ Res. 2016;118(4):620–36. Epub 2016/02/20. doi: 10.1161/CIRCRESAHA.115.306301. PubMed PMID: 26892962; PMCID: PMC4762052.

33. Travaglini KJ, Nabhan AN, Penland L, Sinha R, Gillich A, Sit RV, Chang S, Conley SD, Mori Y, Seita J, Berry GJ, Shrager JB, Metzger RJ, Kuo CS, Neff N, Weissman IL, Quake SR, Krasnow MA. A molecular cell atlas of the human lung from single-cell RNA sequencing. Nature. 2020;587(7835):619-25. Epub 2020/11/20. doi: 10.1038/s41586-020-2922-4. PubMed PMID: 33208946; PMCID: PMC7704697.

34. Kuleshov MV, Jones MR, Rouillard AD, Fernandez NF, Duan Q, Wang Z, Koplev S, Jenkins SL, Jagodnik KM, Lachmann A, McDermott MG, Monteiro CD, Gundersen GW, Ma’ayan A. Enrichr: a comprehensive gene set enrichment analysis web server 2016 update. Nucleic Acids Res. 2016;44(W1):W90–7. Epub 2016/05/05. doi: 10.1093/nar/gkw377. PubMed PMID: 27141961; PMCID: PMC4987924.

35. Kwon AT, Arenillas DJ, Worsley Hunt R, Wasserman WW. oPOSSUM-3: advanced analysis of regulatory motif over-representation across genes or ChIP-Seq datasets. G3 (Bethesda). 2012;2(9):987-1002. Epub 2012/09/14. doi: 10.1534/g3.112.003202. PubMed PMID: 22973536; PMCID: PMC3429929.

36. Han H, Cho JW, Lee S, Yun A, Kim H, Bae D, Yang S, Kim CY, Lee M, Kim E, Lee S, Kang B, Jeong D, Kim Y, Jeon HN, Jung H, Nam S, Chung M, Kim JH, Lee I. TRRUST v2: an expanded reference database of human and mouse transcriptional regulatory interactions. Nucleic Acids Res. 2018;46(D1):D380–D6. Epub 2017/11/01. doi: 10.1093/nar/gkx1013. PubMed PMID: 29087512; PMCID: PMC5753191.

37. Wang Z, Civelek M, Miller CL, Sheffield NC, Guertin MJ, Zang C. BART: a transcription factor prediction tool with query gene sets or epigenomic profiles. Bioinformatics. 2018;34(16):2867–9. Epub 2018/04/03. doi: 10.1093/bioinformatics/bty194. PubMed PMID: 29608647; PMCID: PMC6084568.

38. Looney AP, Han R, Stawski L, Marden G, Iwamoto M, Trojanowska M. Synergistic Role of Endothelial ERG and FLI1 in Mediating Pulmonary Vascular Homeostasis. Am J Respir Cell Mol Biol. 2017;57(1):121–31. Epub 2017/03/02. doi: 10.1165/rcmb.2016-0200OC. PubMed PMID: 28248553; PMCID: PMC5516275.

39. Shah AV, Birdsey GM, Peghaire C, Pitulescu ME, Dufton NP, Yang Y, Weinberg I, Osuna Almagro L, Payne L, Mason JC, Gerhardt H, Adams RH, Randi AM. The endothelial transcription factor ERG mediates Angiopoietin-1-dependent control of Notch signalling and vascular stability. Nat Commun. 2017;8:16002. Epub 2017/07/12. doi: 10.1038/ncomms16002. PubMed PMID: 28695891; PMCID: PMC5508205.

40. Plumitallo S, Ruiz-Llorente L, Langa C, Morini J, Babini G, Cappelletti D, Scelsi L, Greco A, Danesino C, Bernabeu C, Olivieri C. Functional analysis of a novel ENG variant in a patient with hereditary hemorrhagic telangiectasia (HHT) identifies a new Sp1 binding-site. Gene. 2018;647:85–92. Epub 2018/01/07. doi: 10.1016/j.gene.2018.01.007. PubMed PMID: 29305977.

41. Subramanian A, Narayan R, Corsello SM, Peck DD, Natoli TE, Lu X, Gould J, Davis JF, Tubelli AA, Asiedu JK, Lahr DL, Hirschman JE, Liu Z, Donahue M, Julian B, Khan M, Wadden D, Smith IC, Lam D, Liberzon A, Toder C, Bagul M, Orzechowski M, Enache OM, Piccioni F, Johnson SA, Lyons NJ, Berger AH, Shamji AF, Brooks AN, Vrcic A, Flynn C, Rosains J, Takeda DY, Hu R, Davison D, Lamb J, Ardlie K, Hogstrom L, Greenside P, Gray NS, Clemons PA, Silver S, Wu X, Zhao WN, Read-Button W, Wu X, Haggarty SJ, Ronco LV, Boehm JS, Schreiber SL, Doench JG, Bittker JA, Root DE, Wong B, Golub TR. A Next Generation Connectivity Map: L1000 Platform and the First 1,000,000 Profiles. Cell. 2017;171(6):1437–52 e17. Epub 2017/12/02. doi: 10.1016/j.cell.2017.10.049. PubMed PMID: 29195078; PMCID: PMC5990023.

42. Van Quickelberghe E, De Sutter D, van Loo G, Eyckerman S, Gevaert K. A protein-protein interaction map of the TNF-induced NF-κB signal transduction pathway. Scientific Data. 2018;5(1). doi: 10.1038/sdata.2018.289.

43. Ochoa D, Jarnuczak AF, Vieitez C, Gehre M, Soucheray M, Mateus A, Kleefeldt AA, Hill A, Garcia-Alonso L, Stein F, Krogan NJ, Savitski MM, Swaney DL, Vizcaino JA, Noh KM, Beltrao P. The functional landscape of the human phosphoproteome. Nat Biotechnol. 2020;38(3):365–73. Epub 2019/12/11. doi: 10.1038/s41587-019-0344-3. PubMed PMID: 31819260; PMCID: PMC7100915.

44. Neet K, Hunter T. Vertebrate non-receptor protein-tyrosine kinase families. Genes Cells. 1996;1(2):147–69. Epub 1996/02/01. doi: 10.1046/j.1365-2443.1996.d01-234.x. PubMed PMID: 9140060.

45. Shah NH, Lobel M, Weiss A, Kuriyan J. Fine-tuning of substrate preferences of the Src-family kinase Lck revealed through a high-throughput specificity screen. Elife. 2018;7. doi: 10.7554/eLife.35190. PubMed PMID: 29547119; PMCID: PMC5889215.

46. Courtney AH, Lo WL, Weiss A. TCR Signaling: Mechanisms of Initiation and Propagation. Trends Biochem Sci. 2018;43(2):108–23. doi: 10.1016/j.tibs.2017.11.008. PubMed PMID: 29269020; PMCID: PMC5801066.

47. Chakraborty AK, Weiss A. Insights into the initiation of TCR signaling. Nat Immunol. 2014;15(9):798–807. doi: 10.1038/ni.2940. PubMed PMID: 25137454; PMCID: PMC5226627.

48. Hurst LA, Dunmore BJ, Long L, Crosby A, Al-Lamki R, Deighton J, Southwood M, Yang X, Nikolic MZ, Herrera B, Inman GJ, Bradley JR, Rana AA, Upton PD, Morrell NW. TNFalpha drives pulmonary arterial hypertension by suppressing the BMP type-II receptor and altering NOTCH signalling. Nat Commun. 2017;8:14079. doi: 10.1038/ncomms14079. PubMed PMID: 28084316; PMCID: PMC5241886.

49. Betapudi V, Shukla M, Alluri R, Merkulov S, McCrae KR. Novel role for p56/Lck in regulation of endothelial cell survival and angiogenesis. FASEB J. 2016;30(10):3515–26. doi: 10.1096/fj.201500040. PubMed PMID: 27402674; PMCID: PMC5024698.

50. Herazo-Maya JD, Noth I, Duncan SR, Kim S, Ma SF, Tseng GC, Feingold E, Juan-Guardela BM, Richards TJ, Lussier Y, Huang Y, Vij R, Lindell KO, Xue J, Gibson KF, Shapiro SD, Garcia JG, Kaminski N. Peripheral blood mononuclear cell gene expression profiles predict poor outcome in idiopathic pulmonary fibrosis. Sci Transl Med. 2013;5(205):205ra136. doi: 10.1126/scitranslmed.3005964. PubMed PMID: 24089408; PMCID: PMC4175518.

51. Matache C, Onu A, Stefanescu M, Tanaseanu S, Dragomir C, Dolganiuc A, Szegli G. Dysregulation of p56lck inase in patients with systemic lupus erythematosus. Autoimmunity. 2001;34(1):27–38.

52. Pertel T, Zhu D, Panettieri RA, Yamaguchi N, Emala CW, Hirshman CA. Expression and muscarinic receptor coupling of Lyn kinase in cultured human airway smooth muscle cells. American Journal of Physiology-Lung Cellular and Molecular Physiology. 2006;290(3):L492–L500. doi: 10.1152/ajplung.00344.2005.

53. Christmann-Franck S, van Westen GJ, Papadatos G, Beltran Escudie F, Roberts A, Overington JP, Domine D. Unprecedently Large-Scale Kinase Inhibitor Set Enabling the Accurate Prediction of Compound- Kinase Activities: A Way toward Selective Promiscuity by Design? J Chem Inf Model. 2016;56(9):1654–75. Epub 2016/08/03. doi: 10.1021/acs.jcim.6b00122. PubMed PMID: 27482722; PMCID: PMC5039764.

54. Musa A, Ghoraie LS, Zhang SD, Glazko G, Yli-Harja O, Dehmer M, Haibe-Kains B, Emmert-Streib F. A review of connectivity map and computational approaches in pharmacogenomics. Brief Bioinform. 2018;19(3):506–23. Epub 2017/01/11. doi: 10.1093/bib/bbw112. PubMed PMID: 28069634; PMCID: PMC5952941.

55. Gu M, Donato M, Guo M, Wary N, Miao Y, Mao S, Saito T, Otsuki S, Wang L, Harper RL, Sa S, Khatri P, Rabinovitch M. iPSC-endothelial cell phenotypic drug screening and in silico analyses identify tyrphostin- AG1296 for pulmonary arterial hypertension. Sci Transl Med. 2021;13(592). Epub 2021/05/07. doi: 10.1126/scitranslmed.aba6480. PubMed PMID: 33952674; PMCID: PMC8762958.

56. Bai L, Scott MKD, Steinberg E, Kalesinskas L, Habtezion A, Shah NH, Khatri P. Computational drug repositioning of atorvastatin for ulcerative colitis. Journal of the American Medical Informatics Association. 2021;28(11):2325–35. doi: 10.1093/jamia/ocab165.

57. Bhide RS, Cai ZW, Zhang YZ, Qian L, Wei D, Barbosa S, Lombardo LJ, Borzilleri RM, Zheng X, Wu LI, Barrish JC, Kim SH, Leavitt K, Mathur A, Leith L, Chao S, Wautlet B, Mortillo S, Jeyaseelan R, Sr., Kukral D, Hunt JT, Kamath A, Fura A, Vyas V, Marathe P, D’Arienzo C, Derbin G, Fargnoli J. Discovery and preclinical studies of (R)-1-(4-(4-fluoro-2-methyl-1H-indol-5-yloxy)-5- methylpyrrolo[2,1-f][1,2,4]triazin-6-yloxy)propan- 2-ol (BMS-540215), an in vivo active potent VEGFR-2 inhibitor. J Med Chem. 2006;49(7):2143–6. Epub 2006/03/31. doi: 10.1021/jm051106d. PubMed PMID: 16570908.

58. Qi Chao, Kelly G Sprankle, Robert M Grotzfeld, Andiliy G Lai, Todd A Carter, Anne Marie Velasco RNG, Merryl D Cramer, Michael F Gardner, Joyce James, Patrick P Zarrinkar, Hitesh K Patel, Bhagwat SS. Identification of N-(5-tert-butyl-isoxazol-3-yl)-N’-{4-[7-(2-morpholin-4-yl-ethoxy)imidazo[2,1- b][1,3]benzothiazol-2-yl]phenyl}urea dihydrochloride (AC220), a uniquely potent, selective, and efficacious FMS-like tyrosine kinase-3 (FLT3) inhibitor. J Med Chem. 2009;52(23):7808–16. doi: 10.1021/jm9007533.

59. Wilhelm SM, Dumas J, Adnane L, Lynch M, Carter CA, Schutz G, Thierauch KH, Zopf D. Regorafenib (BAY 73-4506): a new oral multikinase inhibitor of angiogenic, stromal and oncogenic receptor tyrosine kinases with potent preclinical antitumor activity. Int J Cancer. 2011;129(1):245–55. Epub 2010/12/21. doi: 10.1002/ijc.25864. PubMed PMID: 21170960.

60. Spiekerkoetter E, Tian X, Cai J, Hopper RK, Sudheendra D, Li CG, El-Bizri N, Sawada H, Haghighat R, Chan R, Haghighat L, de Jesus Perez V, Wang L, Reddy S, Zhao M, Bernstein D, Solow-Cordero DE, Beachy PA, Wandless TJ, Ten Dijke P, Rabinovitch M. FK506 activates BMPR2, rescues endothelial dysfunction, and reverses pulmonary hypertension. J Clin Invest. 2013;123(8):3600–13. doi: 10.1172/JCI65592. PubMed PMID: 23867624; PMCID: PMC3726153.

61. Chen PI, Cao A, Miyagawa K, Tojais NF, Hennigs JK, Li CG, Sweeney NM, Inglis AS, Wang L, Li D, Ye M, Feldman BJ, Rabinovitch M. Amphetamines promote mitochondrial dysfunction and DNA damage in pulmonary hypertension. JCI Insight. 2017;2(2):e90427. Epub 2017/02/01. doi: 10.1172/jci.insight.90427. PubMed PMID: 28138562; PMCID: PMC5256132.

62. Carpentier G, Berndt S, Ferratge S, Rasband W, Cuendet M, Uzan G, Albanese P. Angiogenesis Analyzer for ImageJ - A comparative morphometric analysis of “Endothelial Tube Formation Assay” and “Fibrin Bead Assay”. Sci Rep. 2020;10(1):11568. Epub 2020/07/16. doi: 10.1038/s41598-020-67289-8. PubMed PMID: 32665552; PMCID: PMC7360583.

63. Dobin A, Davis CA, Schlesinger F, Drenkow J, Zaleski C, Jha S, Batut P, Chaisson M, Gingeras TR. STAR: ultrafast universal RNA-seq aligner. Bioinformatics. 2013;29(1):15–21. Epub 2012/10/30. doi: 10.1093/bioinformatics/bts635. PubMed PMID: 23104886; PMCID: PMC3530905.

64. Love MI, Huber W, Anders S. Moderated estimation of fold change and dispersion for RNA-seq data with DESeq2. Genome Biol. 2014;15(12):550. Epub 2014/12/18. doi: 10.1186/s13059-014-0550-8. PubMed PMID: 25516281; PMCID: PMC4302049.

65. Chen EY, Tan CM, Kou Y, Duan Q, Wang Z, Meirelles GV, Clark NR, Ma’ayan A. Enrichr: interactive and collaborative HTML5 gene list enrichment analysis tool. BMC Bioinformatics. 2013;14:128. Epub 2013/04/17. doi: 10.1186/1471-2105-14-128. PubMed PMID: 23586463; PMCID: PMC3637064.

66. Stuart T, Butler A, Hoffman P, Hafemeister C, Papalexi E, Mauck WM, 3rd, Hao Y, Stoeckius M, Smibert P, Satija R. Comprehensive Integration of Single-Cell Data. Cell. 2019;177(7):1888–902 e21. Epub 2019/06/11. doi: 10.1016/j.cell.2019.05.031. PubMed PMID: 31178118; PMCID: PMC6687398.

67. Wang F, Flanagan J, Su N, Wang LC, Bui S, Nielson A, Wu X, Vo HT, Ma XJ, Luo Y. RNAscope: a novel in situ RNA analysis platform for formalin-fixed, paraffin-embedded tissues. J Mol Diagn. 2012;14(1):22–9. Epub 2011/12/15. doi: 10.1016/j.jmoldx.2011.08.002. PubMed PMID: 22166544; PMCID: PMC3338343.

